# Hyaluronan-NK cell Interaction Controls the Primary Vascular Barrier during Early Pregnancy

**DOI:** 10.1101/2020.02.09.940544

**Authors:** Ron Hadas, Eran Gershon, Aviad Cohen, Sima Stroganov, Ofir Atrakchi, Shlomi Lazar, Ofra Golani, Bareket Dassa, Michal Elbaz, Gadi Cohen, Elena Kartvelishvily, Raya Eilam, Nava Dekel, Michal Neeman

## Abstract

Successful implantation is associated with a unique spatial pattern of vascular remodeling, characterized by profound peripheral neo-vascularization surrounding a peri-embryo avascular niche. We hypothesized that hyaluronan controls the formation of the unique vascular pattern encompassing the embryo. This hypothesis was evaluated by genetic modification of hyaluronan metabolism specifically targeted to embryonic trophoblast cells. The outcome of altered hyaluronan deposition on uterine vascular remodeling and post-implantation development were analyzed by MRI, detailed histological examinations, and RNA-sequencing of uterine NK cells. Our experiments revealed that eliminating the anti-angiogenic hyaluronan, led to elevated expression of MMP-9, VEGF-A and its receptor VEGFR-2, accompanied by reduced recruitment of uterine NK cells. Further local decrease in VEGFR-3 resulted in impaired formation of vascular sinuous folds, ectopic angiogenesis and dysfunctional uterine NK cells. Conversely, enhanced deposition of hyaluronan caused the expansion of the maternal-embryo barrier, leading to an increased diffusion distance and aborted implantation. These results demonstrate a pivotal role for hyaluronan in successful pregnancy by fine-tuning the peri-embryo avascular niche and maternal vascular morphogenesis.

## Introduction

The birth of a properly developed mammalian offspring requires the fulfillment of a series of complex, highly regulated processes, initiated by embryo implantation and placental formation concomitant with uterine decidualization, followed by proper fetal development culminating by parturition. A timely and coordinated sequence of these events is subjected to regulation by ovarian steroid hormones, estrogen and progesterone^1^. In humans, the success of natural conception per menstrual cycle is low (∼30%), and implantation defects were implicated in 75% of failed pregnancies ^2 3^.

The general principles of implantation are well conserved among mammals. Implantation occurs at the blastocyst stage, at which the first cell lineages form ^4^. The blastocyst contains an inner cell mass, which mainly gives rise to the fetal organs, and the trophectoderm, an outer epithelial-like cell layer, which attaches to the uterine epithelium. The trophectoderm invades, in turn, the underlying stroma to form the extra-embryonic ectoderm and the ecto-placental cone ^5^, further developing into the hemo-chorial placentae ^6, 7^.

Implantation in mice, which takes place at embryonic day 4.5 (E4.5) starts with blastocyst apposition and its attachment to the uterine epithelium, to trigger decidualization of the uterine stromal cells, characterized by rapid cellular proliferation and differentiation. Secretion of VEGF-A by the decidual cells and induction of VEGF-VEGFR-2 signaling ^8, 9^, result in an immediate local increase in vascular permeability followed by the expression of CD34, an endothelial marker for angiogenesis ^10^. This neo-vascularization event, takes place in the anti-mesometrial pole of the decidua, the site of initial trophoblast invasion and is governed by the progesterone-progesterone receptor axis^8^. The newly formed vessels continue to increase in number and diameter, to supply the embryo with oxygen and nutrients, necessary for its survival and rapid growth ^11^. The decidual mesometrial pole, on the other hand, is characterized by the development of vascular sinuous folds (VSFs), which are arterio-venous vascular shunts, markedly enlarged and elongated prior to placentation ^8^. Another receptor for VEGF signaling, VEGFR-3, participates in this orchestrated series of events as an attenuator of vascular permeability by inhibiting VEGFR-2 expression, consequently blocking VEGF-VEGFR-2 induced permeability ^12, 13^. Perturbation of vascular remodeling during early pregnancy is associated with pathologies, often detected at later stages of gestation, including first-trimester miscarriages, preeclampsia, placental failure and intrauterine growth restriction^8^.

In addition to these strictly controlled angiogenic events, the effective delivery of maternal supply to the embryo is absolutely dependent on the formation of two decidual sub-compartments at E6.5; the highly vascularized secondary decidual zone at the decidual rim, and the avascular primary decidual zone, adjacent to the embryo. This spatially regulated growth of vessels, together with the embryonic diffusion barrier, form the hypoxic niche of the peri-implantation embryo ^14^. This barrier enables maternal nourishing of the embryo while avoiding its immediate exposure the maternal circulation, thus ^15^controlling the delivery of blood-borne high molecular-weight contents, including immune cells and immunoglobulins. ^1, 16^. The mechanism underlying the formation of this diffusion barrier is poorly understood.

Prior to placentation, the decidua, is enriched with a large population of leukocytes, comprised mainly of uterine NK cells, accounting for 70% of the decidual stroma ^17^. Interestingly, in both humans as well as mice, the functions of uterine NK cells differ profoundly from peripheral NK cells. Namely, unlike the predominant cytotoxic actions against virus-infected or cancerous cells, characteristic to peripheral NK cells, the uterine subsets show differential effector repertoire and vessel remodeling activities^15, 18 17^. Moreover, uterine NK cells, have been demonstrated to regulate ‘ligand-independent’ VEGFR-3 coordination of enlargement and elongation of VSFs via pruning of mesometrial vasculature ^8^.

Blastocyst adhesion to the uterine lining, a necessary prelude for implantation, is facilitated by delicate paracrine and juxtacrine interactions. The latter are characterized mainly by extra-cellular matrix (ECM) components, including specific mucins, selectins, integrins, and glycosaminoglycans, such as heparan sulfate and hyaluronan ^2^. During trophoblast invasion, uterine epithelial cells locally undergo apoptosis and are phagocytosed by primary trophoblast giant cells. The subsequent basement membrane degradation enables the establishment of direct contacts between invading trophoblast cells and the endometrial stroma, including uterine decidual cells, decidual blood vessels and their supporting ECM ^19^. Following trophectoderm invasion, of the anti-mesometrial pole of the uterine lumen ^19^, the ECM undergoes extensive transformations, including generation of an interrupted and thickened basement membrane within the peri-cellular spaces of the post-implantation decidua. The latter may confine infiltration of immune cells, or block exposure to maternal antibodies, to sin the frame of primary maternal tolerance ^20^. Trophoblast invasion is facilitated by trophectoderm cells-produced proteinases, such as MMP-9, and urokinase-type plasminogen activator ^2^, that modify the ECM and possibly exploit uterine ECM remodeling, including rapid endometrial hyaluronan clearance ^21^.

Hyaluronan is a negatively charged un-branched cell-surface associated polysaccharide, composed solely of repeating disaccharides units of [D-glucuronic acid-β1,3-N-acetyl-D-glucosamine-β1,4-]_n_ ^22^. Hyaluronan synthesis is catalyzed by three hyaluronan synthases, HAS-1, HAS-2 and HAS-3, all anchored to the plasma membrane, where the synthesis occurs. The hyaluronan produced is extruded to the ECM, in its intact polysaccharide form in a typical molecular mass of 10^6^ to 10^7^ Da ^23^. Hyaluronan, in its high molecular-weight form, inhibits angiogenesis in a size-dependent manner *in vitro* ^24^ and *in vivo* ^24–27^. Hyaluronan is cleaved by hyaluronidases. In both human and mouse, there are six hyaluronidase genes encoding for enzymes with different enzymatic properties and cellular localizations ^22, 28^. The main enzymes studied are Hyal-1 and Hyal-2. The most abundant plasma and urine enzyme Hyal-1, intra-cellularly degrades hyaluronan to small oligosaccharides ^23^. Hyal-2, which is anchored to the plasma membrane by a glycosylphosphatidylinositol link, hydrolyzes only high molecular-weight hyaluronan ^23^, and generates intermediate hyaluronan fragments of approximately 20 kDa. These fragments are characterized by their pro-angiogenic activity, including induction of endothelial cell proliferation and tube formation, by mechanisms involving VEGF release and up-regulation of VEGFR-2 expression ^28–30 24, 27, 31^. Hyaluronan oligosaccharides are also potent inducers of MMP-9 activity ^32, 33^ and stimulators of pro-inflammatory macrophages ^23, 34 35, 36^. MMP-9 can independently induce angiogenesis by releasing the ECM-bound VEGF, potentially increasing its abundance near VEGFR-2 expressing cells, as previously demonstrated during carcinogenesis ^37^.

In the study presented here, the role of hyaluronan as a vascular morphogen shaping the primary maternal-embryo barrier was evaluated during implantation and early mouse pregnancy. To this end, we employed lentivirus-mediated genetic modification of hyaluronan metabolism, directed at the embryonic trophectoderm. We report herein that disruption of hyaluronan synthesis, as well as its increased cleavage, at the embryonic niche, impairs implantation by induction of decidual vascular permeability, defective VSF formation and breach of the maternal-embryo barrier. This phenotype was associated with elevated MMP-9 expression and interrupted uterine NK cells recruitment and function. Conversely, enhanced deposition of hyaluronan resulted in the expansion of the maternal-embryo barrier and increased diffusion distance, leading to compromised implantation. We also demonstrate that deposition of hyaluronan at the embryonic niche is subjected to regulation of the PR. These results demonstrate a pivotal role for hyaluronan in the success of pregnancy by fine-tuning the peri-embryo vascular morphogenesis, so as to maintain the primary maternal-embryo vascular barrier and the hypoxic avascular niche, during early pregnancy.

## Results

### Decidual angiogenesis mirrors spatially and temporally the pattern of deposition of hyaluronan and its degradation products

Following embryo implantation, the uterus undergoes rapid and profound tissue remodeling (**Figure 1A**). The previously reported induction of angiogenesis ^10^ is herein indicated by CD34, a marker for newly formed blood vessels observed in the anti-mesometrial pole of the decidua (**Figure 1B**). This vascular modification coincided spatially and temporally with dynamic alterations in the deposition of hyaluronan after implantation. Specifically, following implantation (E5.5), hyaluronan accumulated in the mesometrial poleand surrounding the embryo. At E6.5 hyaluronan was mainly deposited in the ecto-placental cone in mesometrial orientation to the implantation chamber (**Figure 1C**). Accumulation of hyaluronan oligosaccharides was detected in pregnant mice following embryo implantation, in concomitance intensified decidual angiogenesis (**Figure 1D**).

**Figure 1.**
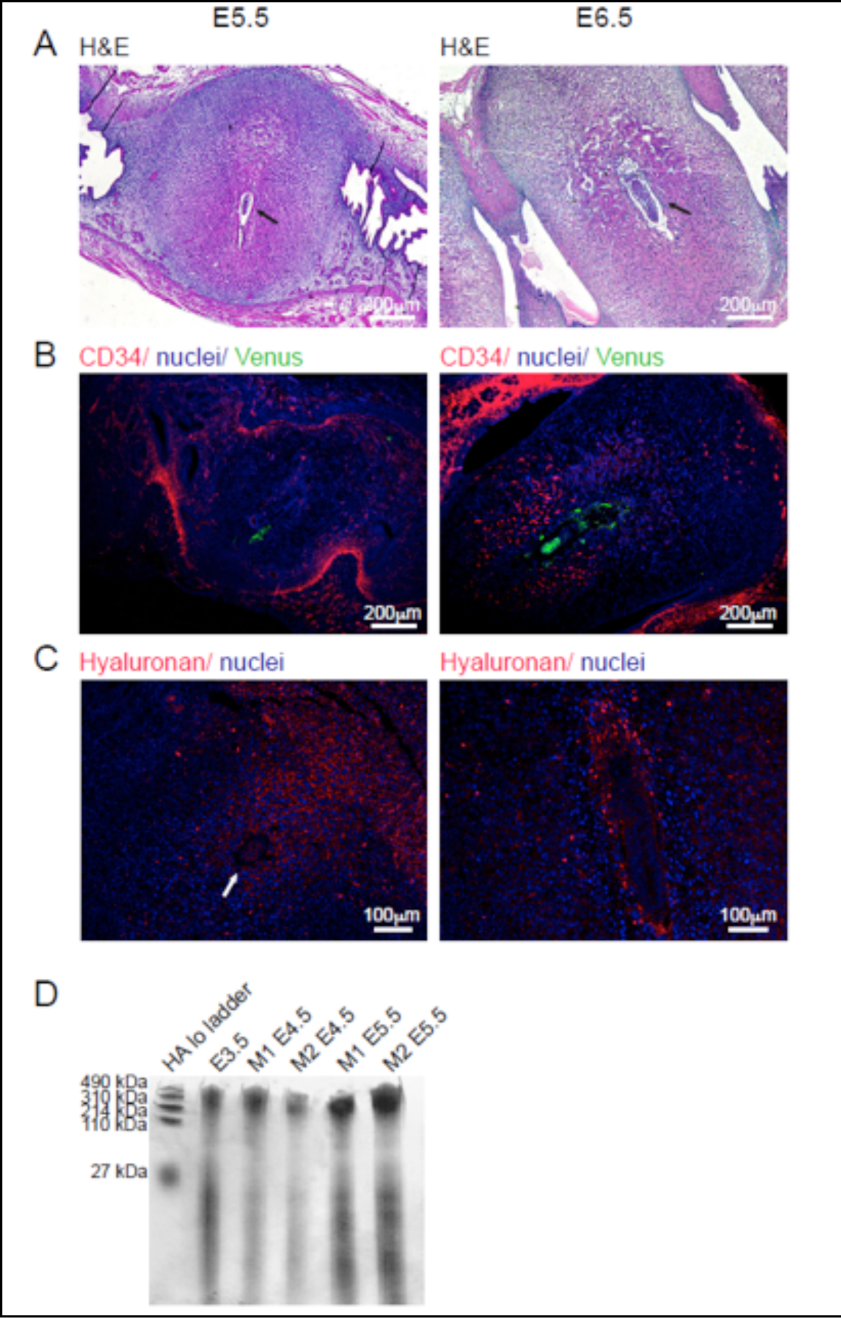
Hyaluronan deposition and vascular remodeling in the implantation site during early pregnancy. Female mice were mated with Venus+ males and their uterine horns were harvested during the peri-implantation period, from E5.5 to E6.5. (n=4 dams). (**A**) H&E staining of deciduae. Black arrows designate embryos. (**B**) Newly formed CD34^+^ uterine blood vessels, reflecting decidual angiogenesis. (**C**) Hyaluronan localization by immunohistochemistry. White arrow designate embryo. (**D**) Equal number of embryo implantation sites were harvested from each mouse. Glycosaminoglycans were separated from uteri of pregnant mice, at different time points, subjected to native PAGE next to Hyaluronan standard and stained for hyaluronan (n=2 dams; 3 implantation sites per dam).

To further explore hyaluronan deposition we analyzed the distribution of hyaluronan synthases, hyaluronidases and hyaluronan binding proteins throughout the peri-implantation period. We found that the hyaluronan synthesizing enzyme, HAS-1 was locally up-regulated in implantation sites at E4.5, transiently decreasing at E5.5, and increasing again at E6.5 (**Figure S2A**). Interestingly, the expression of HAS-2 continuously decreased during the post-implantation period (**Figure S2B**). Hyal-1 expression underwent gradual down-regulation (**Figure S2C**), whereas the expression levels of Hyal-2 remained stable throughout implantation (**Figure S2D**). We also analyzed the expression pattern of two hyalhedrines, hyaluronan ECM stabilizing glycoproteins, TSG-6 and Versican. TSG-6 was significantly up-regulated on the day of implantation, followed by a sharp reduction in its expression (**Figure S2E**), whereas Versican expression levels remained constant during the two consecutive days following blastocyst attachment and invasion (**Figure S2F**).

For further resolution, we examined the spatio-temporal distribution of hyaluronan bio-synthesis enzymes in the different decidual sub-compartments (**Figure 2**). This analysis revealed that post-implantation, at E5.5, HAS-1 was expressed solely by maternal cells in the primary decidual zone, forming a sphere surrounding the embryo, while HAS-2 was expressed both by giant as well as cytotrophoblast cells at the ecto-placental cone. At E6.5, both hyaluronan synthases were distributed in the anti-mesometrial pole of the decidua. HAS-1 was expressed in maternal cells adjacent to trophoblast cells, while HAS-2 alone was detected robustly in the ecto-placental cone at the mesometrial region of the embryonic egg cylinder, and by trophoblast giant cells (**Figure 2A**&**B**).

**Figure 2.**
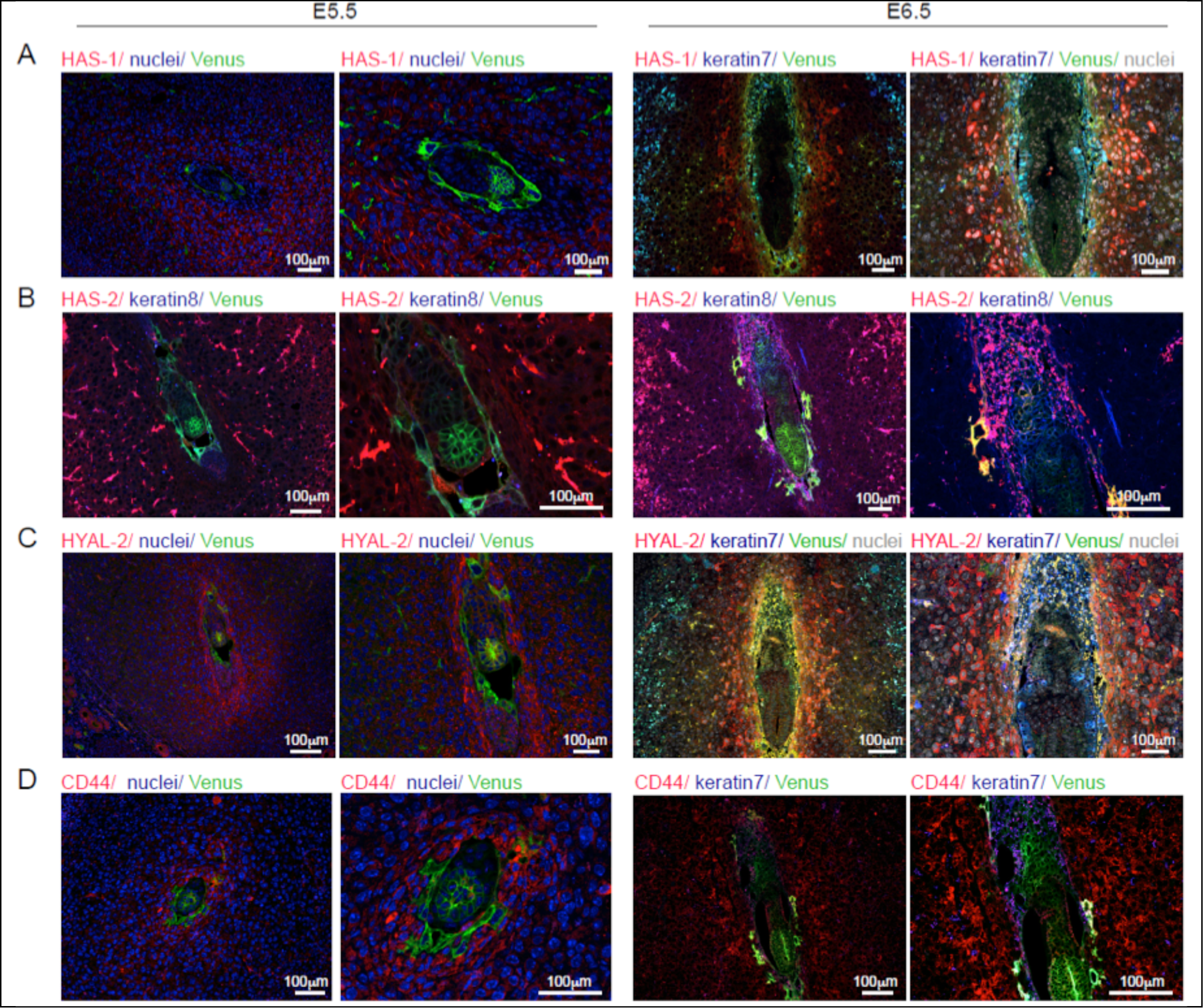
Hyaluronan metabolism following embryo implantation. Female mice were mated with Venus+ males and their uterine horns were harvested during the peri-implantation period, from E5.5 to E6.5 specific histological analysis were conducted alongside Venus (embryo) and keratin that detects trophoblast and visceral endoderm cells. (n=4 dams, 10 implantation sites) left panels are magnifications of right panels. (**A-B**) Immunohistochemical analysis of hyaluronan synthesizing enzymes HAS-1 and HAS-2 (C) Immunohistochemical analysis of Hyal-2, hyaluronan degrading enzyme. (**D**) Immunohistochemical analysis of the most prominent hyaluronan receptor, CD44.

At E5.5 hyaluronan degrading enzymes, Hyal-2 and Hyal-1, were both detected in the maternal primary decidual zone, while Hyal-2 was also expressed by all subtypes of trophoblast cells. At E6.5, hyaluronidases were expressed mainly by trophoblast giant cells and adjacent decidualized cells, and to a lesser extent by embryonic endoderm cells surrounding the embryo, (**Figure 2C**& **Figure S2G**).

Not surprisingly, two of the receptors for hyaluronan oligosaccharides, CD44 and LYVE-1^22^, were co-expressed with HAS-2 and the two Hyaluronidases. After the day of implantation, both CD44 and LYVE-1 showed a pattern of expression similar to that of Hyal-2 and Hyal-1 (**Figure 2D**& **Figure S2H**). Interestingly, RHAMM, a hyaluronan mediated motility receptor, was highly expressed by the maternal decidualized cells, in a ring-like structure at E5.5 and virtually absent at E6.5 (**Figure S2I**).

Overall, we show the close spatial association of hyaluronan bio-synthesis, enzymatic degradation and receptors at the feto-maternal interface post-implantation.

### Progesterone receptor signaling positively regulated hyaluronan degradation, trophoblast invasion and decidual angiogenesis

To examine the involvement of progesterone in hyaluronan metabolism, we suppressed PR signaling via single administration of its antagonist, RU486, at E4.5, post-implantation (8 mg/kg BW). Interestingly, RU486 treatment led to decreased expression of Hyal-2 and impaired anti-mesometrial stromal invasion by primary trophoblast giant cells, at E5.5 (**Figure 3C&D**). As previously demonstrated for later pregnancy^8^, RU486 administration after initial decidualization, resulted in a decreased VEGF-A expression (**Figure 3A**), accompanied by resorptions of embryos (**Figure 3B**).

**Figure 3.**
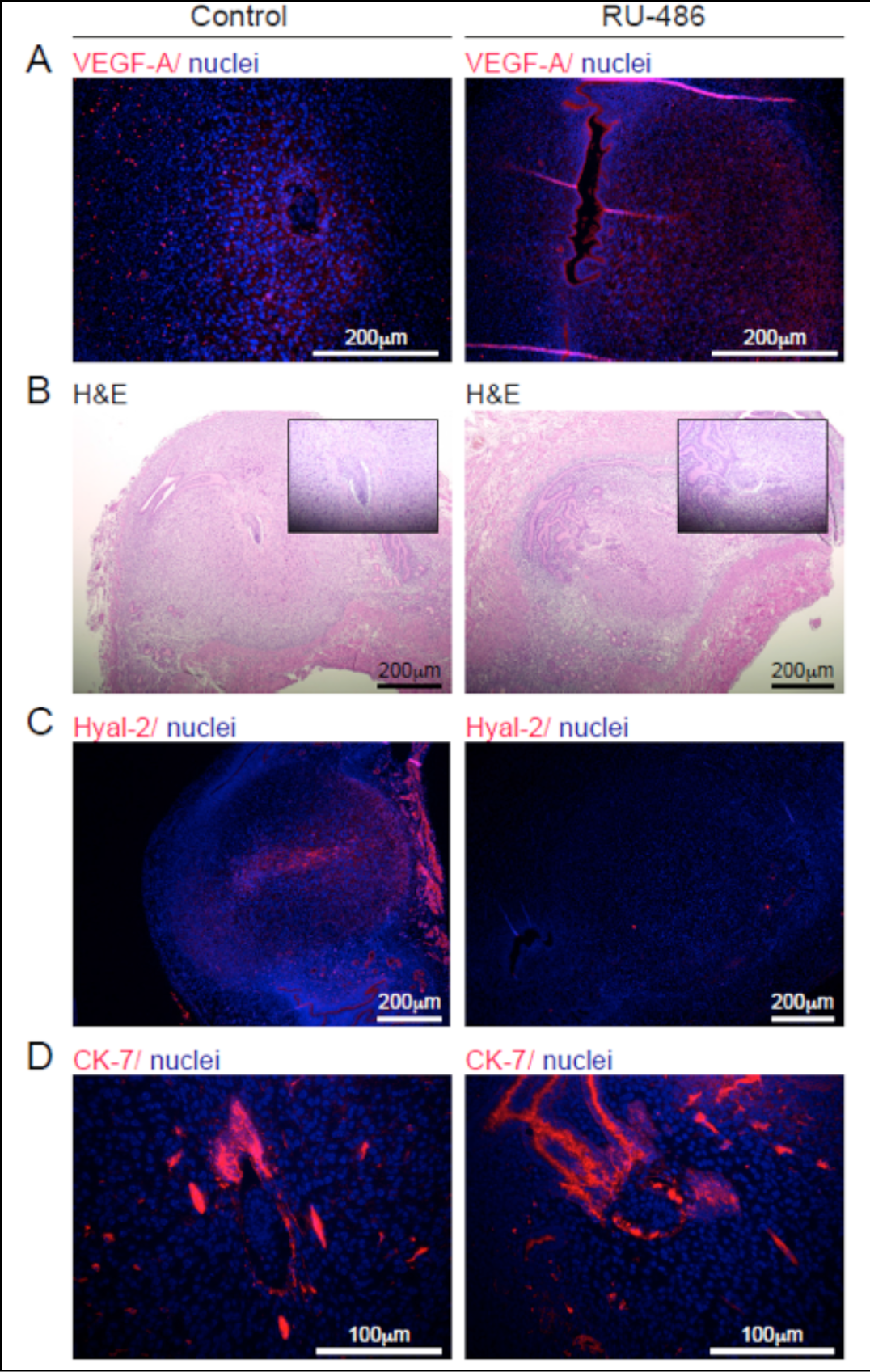
Pharmacological blockade of the progesterone receptor post-attachment. Female ICR mice were mated with ICR males, administered with RU-486 at E4.5, their uterine horns were harvested at E5.5. (n=3 dams). (**A**) Representative images of VEGF-A staining in E5.5 deciduae. (**B**) Smaller deciduae and abnormal embryonic morphology, detected by H&E staining. (**C**) Representative images of decreased Hyal-2 decidual distribution. (**D**) Comparison of Cytokeratin-7 expressing trophoblast cells and their distribution.

### Hyal-2 over-expression in trophoblast cells disrupted the hyaluronan maternal-embryo barrier, leading to embryo resorption

To study the role of hyaluronan at the feto-maternal interface, post-implantation, we generated blastocysts expressing eGFP in their trophectoderm cells along with over-expression of Hyal-2, using lentiviral infection (**Figure 4A**).

**Figure 4.**
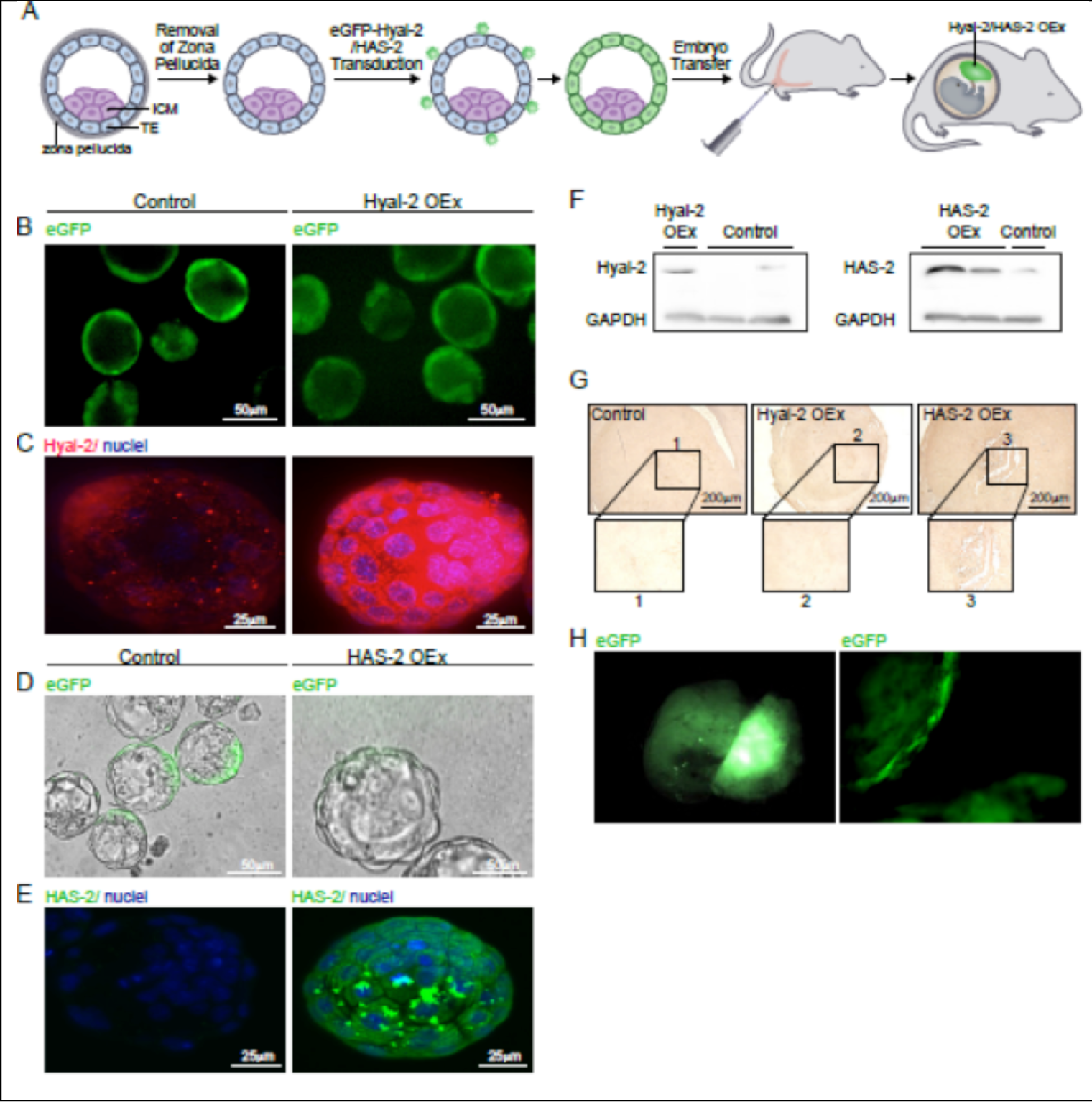
Genetic modifications in embryonic trophoblast cells. Blastocysts over-expressing Hyal-2, HAS-2 and eGFP were generated by lentivirus infection; (**A**) Retrieved from pregnant mice at E2.5, infected with lenti-viral vectors then transferred to pseudo-pregnant mice at E2.5. (**B-C**) Hyal-2 was over-expressed in the trophectoderm. Upper panels show eGFP expression prior to embryo transfer; lower panels show maximal intensity projections of whole-mount immunofluorescence for Hyal-2 in blastocysts following lentiviral transduction (**D-E**) prior to embryo transfer, blastocysts over-expressing HAS-2 in their trophectoderm were visualized for eGFP expression using fluorescent microscopy, and HAS-2 over-expression following viral infection. (**F**) Validation of Hyal-2 and HAS-2 over-expression by Western blot analysis. (**G**) Histological assessment of hyaluronan deposition as a result of Hyal-2 and HAS-2 over-expression. (n=3 mice) (**H**) Placental trophoblast cells expressing eGFP at E12.5 alongside the embryo; trophoblast giant cells expressing eGFP.

Prior to their transfer to pseudo-pregnant recipient mice, the blastocysts were visualized for eGFP expression. Notably, fluorescence was restricted to the trophectoderm, and could not be detected in the ICM (**Figure 4B**&**D**). Transgene over-expression was validated by immunofluorescence of infected embryos (**Figure 4C**&**E**) as well as by Western blot analysis (**Figure 4F**). Hyaluronan was modified as expected following implantation in foster dams (Figure4G). Lentiviral transduction of trophoblasts remained effective following placentation visualized by eGFP visualization (**Figure 4H**).

Over-expression of Hyal-2 in trophoblast cells resulted in augmented mesometrial accumulation of maternal blood (**Figure 5A**). Multiple embryo resorptions and embryonic cell death were observed in pregnant mice carrying Hyal-2 over-expressing (Hyal-2 OEx) blastocysts (**Figure 5B**). Interestingly, each decidua was reduced in diameter, whereas their total number in these mice was significantly higher, compared to controls (**Figure 5C-D**). The transfer of Hyal-2 OEx blastocysts resulted in a robust activation of macrophages, associated with blunt-ended mesometrial blood vessels, demonstrated by staining for MAC-2, a marker for inflammatory macrophages and visualized both by immunohistochemistry and by tissue clearing (**Figure 5E-G**).

**Figure 5.**
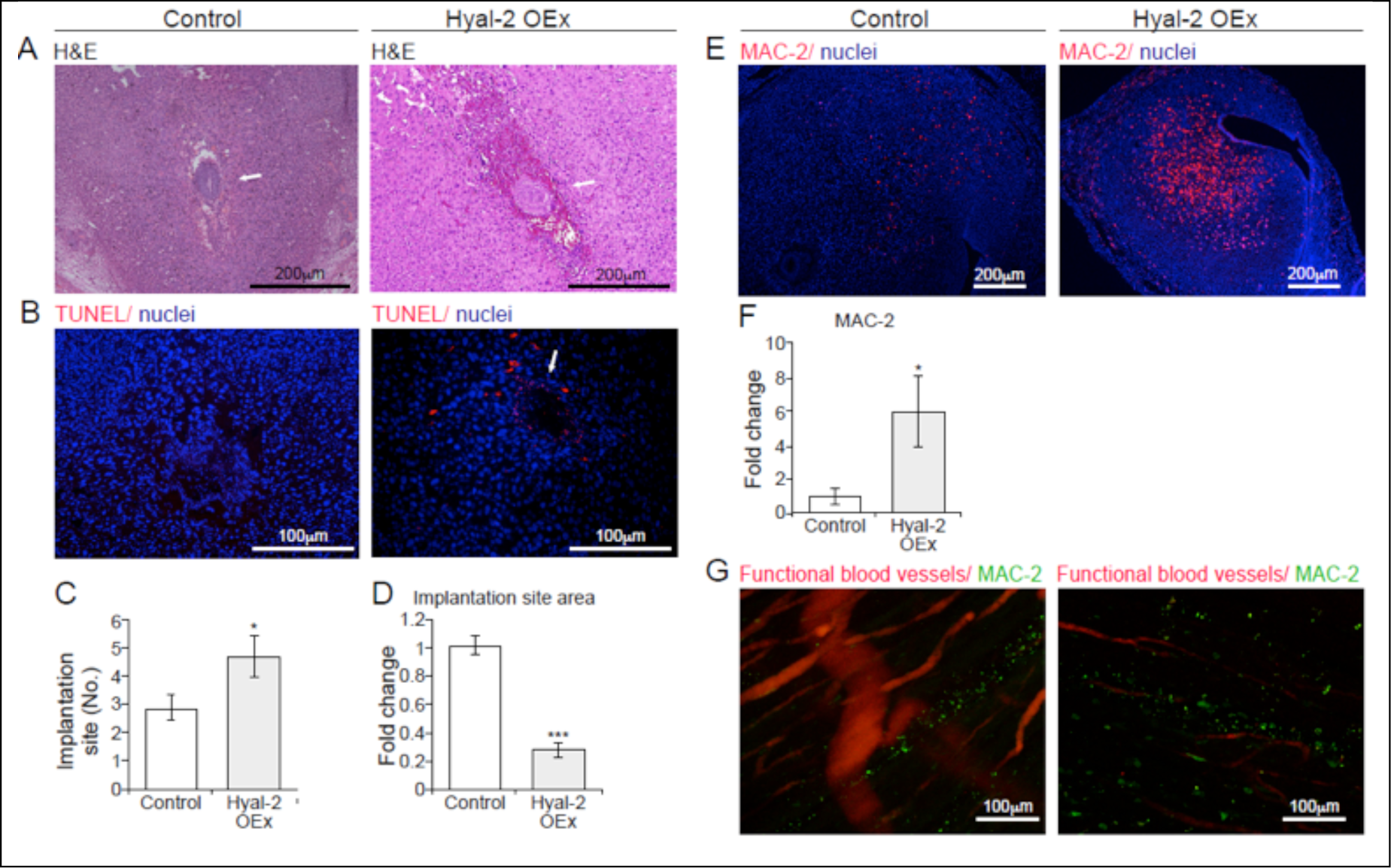
Trophectoderm over-expression of Hyal-2 results enhanced implantation but early embryonic lethality. Implantation sites were studied at E6.5, containing sham infected embryos and embryos over-expressing Hyal-2 in their trophoblast cell layer. (**A**) H&E staining (n=6 dams; white arrows, deciduae). (**B**) TUNEL staining revealed cells undergoing cell death at the embryonic niche of Hyal-2 OEx (white arrow, embryonic cells; n=4 dams). (**C-D**) Quantification of observed deciduae as well as their calculated area. (2.9±0.5; 4.75± 0.7; p=0.02) (n=12 dams in control, n=10 dams in Hyal-2 OEx) /(3.42 fold ±0.07; 0.03; p=0.0006) (n=8 dams, 27 implantation sites in control; n=6 dams, 23 implantation sites for Hyal-2 OEx). (**E-F**) Detection of MAC-2^+^ macrophages by fluorescent microscopy and their quantification (5.9 fold change±0.35; p=0.02) (n=5 dams, 9 deciduae from each group). (**G**) Presence of MAC-2^+^ macrophages was further demonstrated using immuno-staining of macrophages in implantation sites harvested from surrogate mothers injected with ROX labeled lectin. Tissues were made transparent using modified Clarity procedure, thus enabling visualization of whole deciduae by confocal microscopy. (n=2 dams from each group).

Interestingly, Hyal-2 OEx resulted with augmented angiogenesis in the embryonic niche (**Figure 6A**). MRI inspection of pregnant mice carrying Hyal-2 OEx blastocysts detected enhanced accumulation of intravenously injected biotin-BSA-Gd DTPA as early as 3 minutes after its administration (**Figure 6B**). Further analysis revealed a significantly increased blood volume fraction with no change in vessel permeability (**Figure 6C-E**). Hyal-2 OEx in the trophoblasts led to increased levels of VEGF-A, and up-regulation of VEGFR-2, the latter observed specifically in mesometrial orientation to the embryo and the ecto-placental cone (**Figure 6F&Figure 6G-H**). Notably, VEGFR-1 levels did not differ following hyaluronan increased degradation (**Figure S4A**). However trophoblast Hyal-2 OEx, brought about a decreased VEGFR-3 expression in the mesometrial pole. This last effect resulted in diminished VSFs formation (**Figure 6I, J& Figure S4B**), accompanied by mesometrial contrast agent accumulation (**Figure 6J**). Thus, enhanced enzymatic degradation of hyaluronan was sufficient for breaching the primary maternal-embryo vascular barrier and perturbing the spatial balance of decidual vascular remodeling.

**Figure 6.**
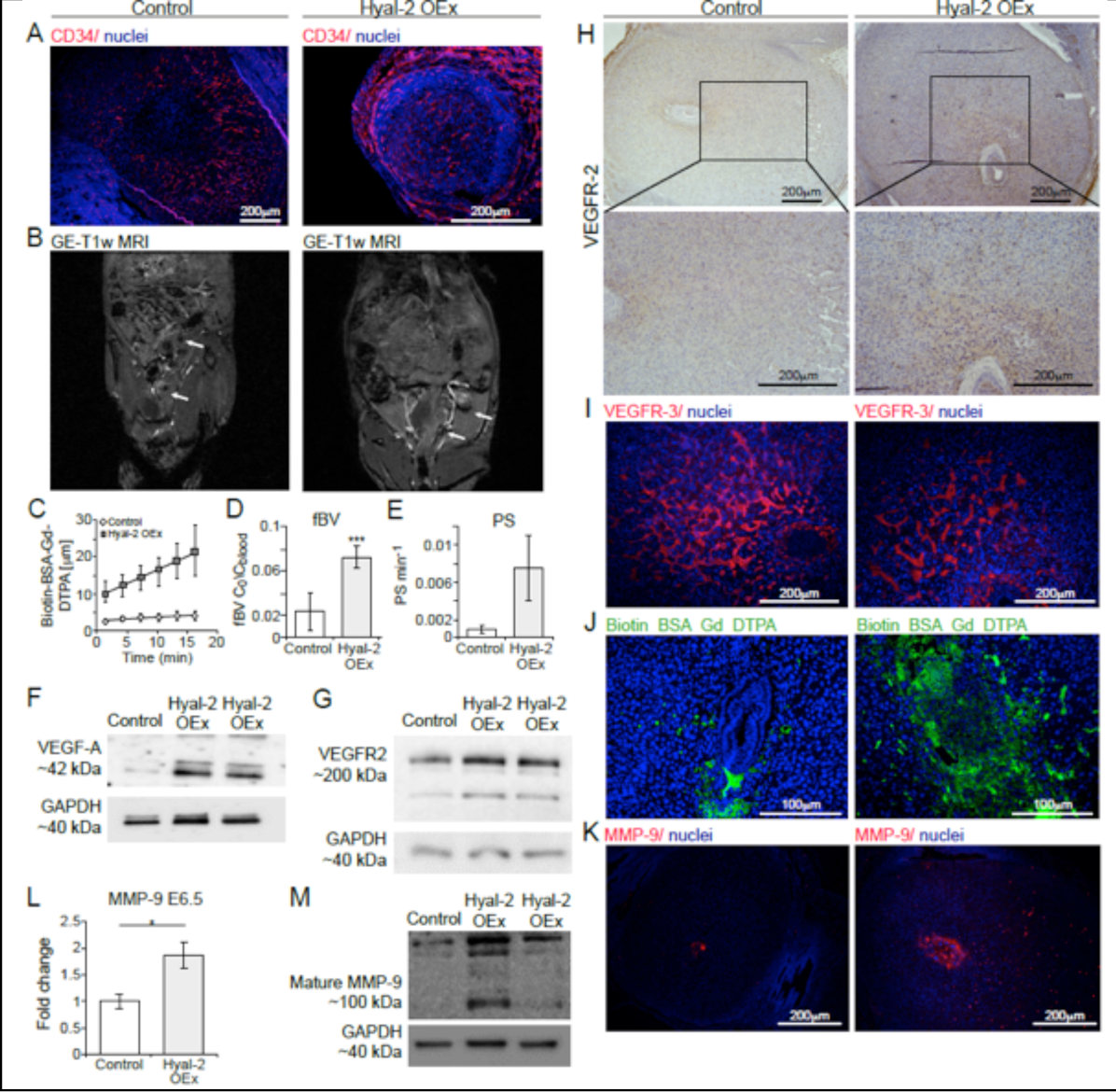
Detrimental decidual hyper-vascularity and breach of the maternal-embryo barrier is induced by trophectoderm Hyal-2 over-expression. (**A**) Ectopic presence of newly formed maternal blood vessels in the embryonic niche was observed in Hyal-2 OEx dams, as opposed to the control group (n=5 dams). (**B**) GE-T1 weighted images of deciduae, acquired from pregnant mice at E6.5, 3 minutes after administration of biotin-BSA-GdDTPA.; white arrows point to implantation sites. (**C-E**) Linear regression plots from DCE MRI analysis; Decidual blood volume fraction (fBV; 0.023±0.01; 0.072±0.009, p=0.0005) and permeability (PS; 0.00092±0.0003; 0.0074±0.0031, p=0.09) were calculated from DCE MRI (n=5 dams 13 implantation sites in control; 6 dams 24 implantation sites in Hyal-2 OEx). (**F-G**) Increased levels of VEGF-A and VEGFR-2 in deciduae harvested at E6.5 (n=3 dams 2 implantation sites from each group). (**H**) Increased VEGFR-2 expression on decidual vessels in mesometrial orientation as well as in cytotrophoblast cells, in Hyal-2 OEx foster dams (n=4 dams in each group). (**I**) Declined VEGFR-3 positive VSFs endothelial expression, demonstrated by immunofluorescence (n=3 dams 6 implantation sites from each group). (**J**) Hyper-permeable blood vessels in the embryonic niche were visualized by staining of biotin-BSA-GdDTPA, 40 minutes after intravenous injection (n=5 dams). (**K**) Increased MMP-9 levels in E6.5 trophoblast cells, as a result of Hyal-2 over-expression (1.858 fold change±0.14; 0.25 p=0.01) (n=5 dams in each group). (**L**) Quantification of MMP-9 expression in E6.5 deciduae by immunofluorescence of histological sections (1.858 fold change±0.14; 0.25 p=0.01) (n=5 dams in each group). (**M**) Increased levels of pro MMP-9 and mature MMP-9 in deciduae harvested at E6.5 (n=3 dams 2 implantation sites from each group).

Interestingly, Hyal-2 OEx resulted in increased MMP-9 expression in trophoblast cells, post-implantation (E6.5). The expression of MMP-9 was examined by immunofluorescence as well as by Western blot analysis (**Figure 6K-M**). As expected, this increased expression was accompanied by excess invasion of trophoblast cells at E6.5 (**Figure S4C-D**).

### Hyal-2 over-expression in trophoblast cells resulted in impaired uterine NK cells recruitment and function

Uterine NK cells were detected in the mesometrial pole of E6.5 decidua and flanking trophoblast giant cells (**Figure 7A**). The latter demonstrated the potential of binding hyaluronan by expressing genes encoding for hyaluronan receptors such as: RHAMM (Hmmr), Lyve1, Cd44, Tlr4 (**Figure S2E**). Alongside the altered decidual vasculature observed in foster dams carrying Hyal-2 OEx embryos, uterine NK cells recruitment was largely impaired, as indicated by the reduction in IL15 -IL15R binding assay (**Figure 7B**). This effect evidently brought about a decrease in the abundance of specialized uterine NK cells (**Figure 7C-E**). The diminished accumulation of the uterine NK cells in the mesometrial pole, and decreased expression of uterine NK cells classification marker TNFRSF-9 (**Figure 7F**) correlated with the impaired VSF enlargement observed above (**Figure 6I**), thereby pointing out the consequences diminished decidual hyaluronan inflicts on uterine NK cells and their vascular remodeling potency.

**Figure 7.**
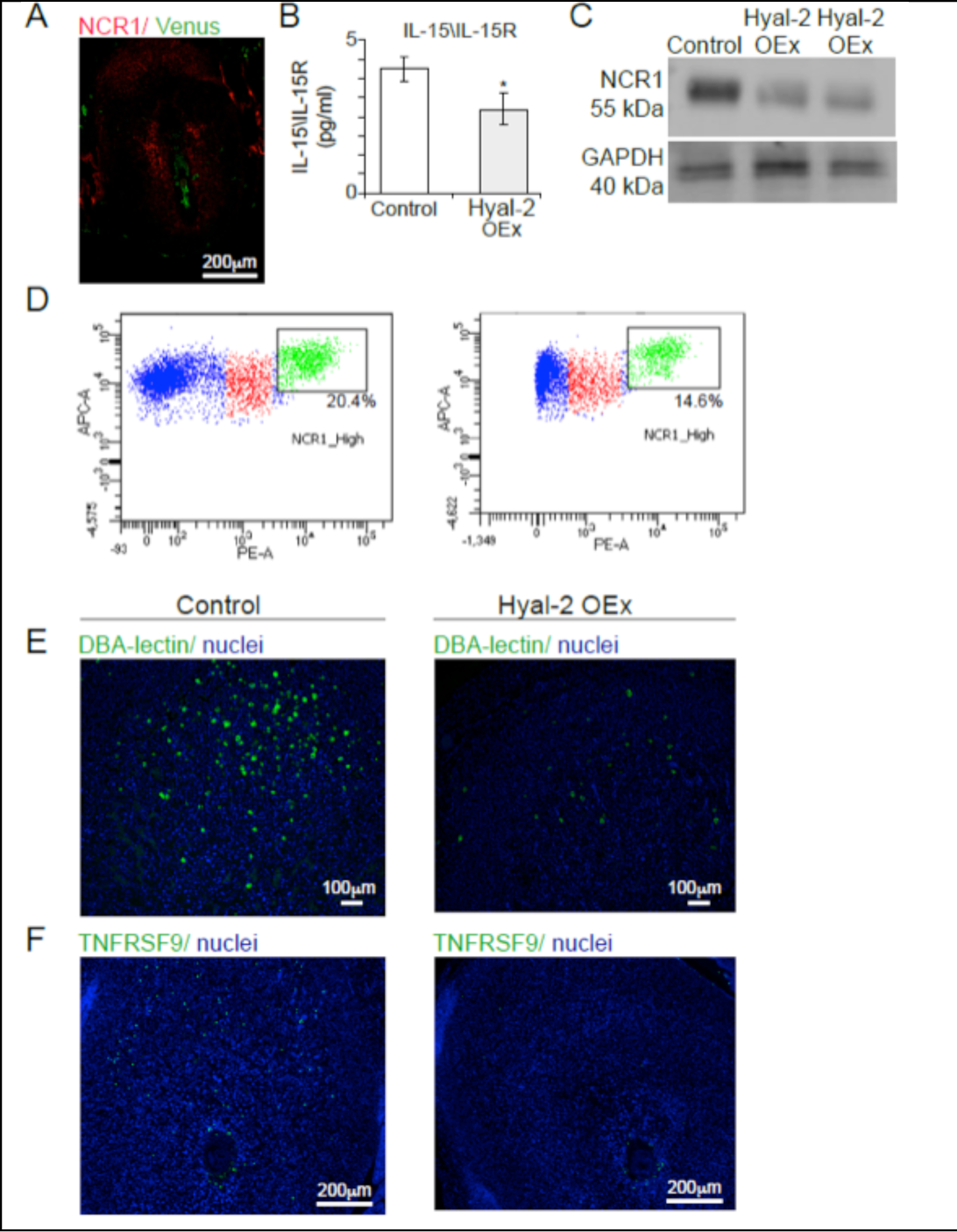
Trophoblast Hyal-2 over-expression impairs uterine NK cells recruitment and function. (**A**)Representative images of NCR-1+ uterine NK cells staining in E6.5 deciduae (n=4 dams in each group). (**B**) Impaired uterine NK cells recruitment demonstrated by decreased IL-15-IL-15R complexes detected by ELISA (n=5 dams in each group 323.4pg/ml±23.49; 213.65pg/ml±30.43, p=0.2) (**C**) Western blot analysis for NCR-1 in E6.5 decidual extracts demonstrated decreased accumulation of NCR-1 expressing NK cells in Hyal-2 OEx foster dams (n=3 dams 2 implantation sites from each group). (**D**) Flow cytometry analysis of E6.5 implantation sites harvested from foster dams of both groups. Note decreased ratio of CD45+NCR-1+ population in the Hyal-2 OEx group. (**E**) Immunofluorescence of DBA+ differentiated uterine NK cells staining in E6.5 deciduae. (**F**) Immunofluorescence of TNFRSF9 expressed by uterine NK cells, in the mesometrial pole in E6.5 deciduae.

### Augmented hyaluronan degradation severely alters uterine NK cells differentiation and homeostatic shift

Transcriptome profiling of NCR-1 expressing uterine NK cells sorted from E6.5 deciduae revealed 46 differentially expressed genes between uterine NK cells sorted from control dams and uterine NK cells sorted from Hyal-2 OEx foster dams. Expression of genes associated with the innate immune response was increased in Hyal-2 OEx uterine NK cells, as opposed to genes associated with unique uterine NK cells classification, differentiation, machinery of protein translation and secretion, and immunomodulation, whose expression was strongly decreased (**Figure 8A**). Functional analysis of sorted uterine NK cells from Hyal-2 OEx dams, revealed significantly reduced protein translation machinery, intra and extra-cellular secretion machinery, vascular remodeling factors and granzyme-mediated apoptotic signaling traditionally associated with uterine NK cells. (Ctsg, Gzm c,e,f,g, Sec61b, Spp1, Tnfrs9) (**Figure 8B&C**). Functionality of uterine NK cells sorted from Hyal-2 OEx dams differed significantly, revealing enrichment of multiple genes, which regulate the inflammatory response and take part in allograft rejection signaling (Tnf, Ccr2, Samd9l, B2m, Npm1, Ly86), diverting from their classification as immunomodulatory NK cells (**Figure 8D&E**).

**Figure 8.**
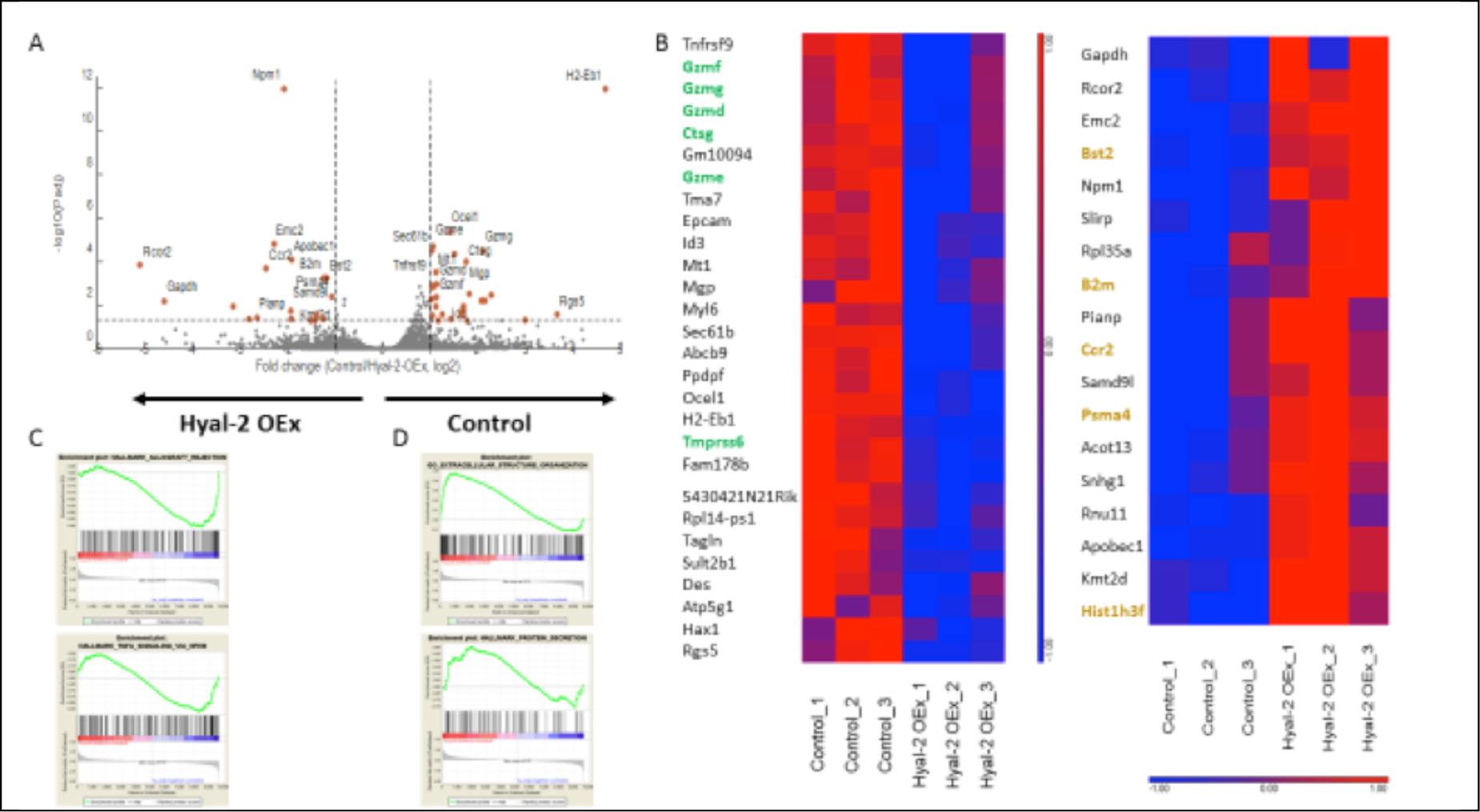
Uterine NK cells transcriptome is dysregulated in Hyal-2 OEx foster dams. NCR1+ uterine NK cells were sorted from E6.5 deciduae of surrogate mice. (n= 3 dams dams; pools of 4 implantation sites each). (**A**) Volcano plot of all genes detected in RNA sequencing analysis. All points above the grey dotted horizontal line are statistically significant. Genes associated with antiviral and innate immune response are indicated in Hyal-2 OEx; Genes associated with uterine NK cells classification are indicated in Control. (**B**) Heat-map of differentially expressed (DE) genes in uterine NK cells sorted from Control dams. Bolt DE genes in green were associated Serine-type Peptidase Activity, characteristic of uterine NK cells by GO: molecular function; Bolt DE genes in orange were associated Cytokine Signaling in Immune System, by GO: Pathways. (**C**) GSEA analysis of uterine NK cells, sorted from Hyal-2 OEx dams, tested for transcriptional enrichment in the above indicated pathways; Genes clustered above the dotted line are significantly enriched. (**D**) GSEA analysis of uterine NK cells, sorted from Control dams, tested for transcriptional enrichment in the above indicated pathways; Genes clustered above the dotted line are significantly enriched.

### HAS-2 over-expression in trophoblast cells resulted in decreased permeability of decidual blood vessels, leading to early embryonic lethality

To further confirm the contribution of hyaluronan metabolism to the success of implantation, lentiviral blastocyst infection was also employed for HAS-2 over-expression (HAS-2 OEx). As expected, the deposition of peri-embryo hyaluronan was enhanced upon the transfer of these blastocysts to pseudo-pregnant mice (**Figure 4**). Nevertheless, while no effect on infiltration of inflammatory macrophages or on the number of implantation sites (**Figure 9D-F**) was observed, these pregnancies resulted in multiple embryo resorptions (**Figure 9A-C**). Furthermore, upon the transfer of HAS-2 over-expressing blastocysts, implantation sites showed impaired formation of new blood vessels (**Figure 9G**) and decreased accumulation of intravenously injected biotin-BSA-Gd DTPA, as detected by MRI 20 minutes after its administration (**Figure 9H**). This decrease was further validated by histological staining of the contrast agent 40 minutes post-injection (**Figure 9I**). Moreover, DCE MRI analysis of pregnant mice carrying HAS-2 OEx embryos, revealed vascular changes opposite to those induced by Hyal-2 OEx (**Figure 9J**), with significantly decreased fBV (**Figure 9K**) and blood-vessel permeability (**Figure 9L**) as compared to control pregnancies.

**Figure 9.**
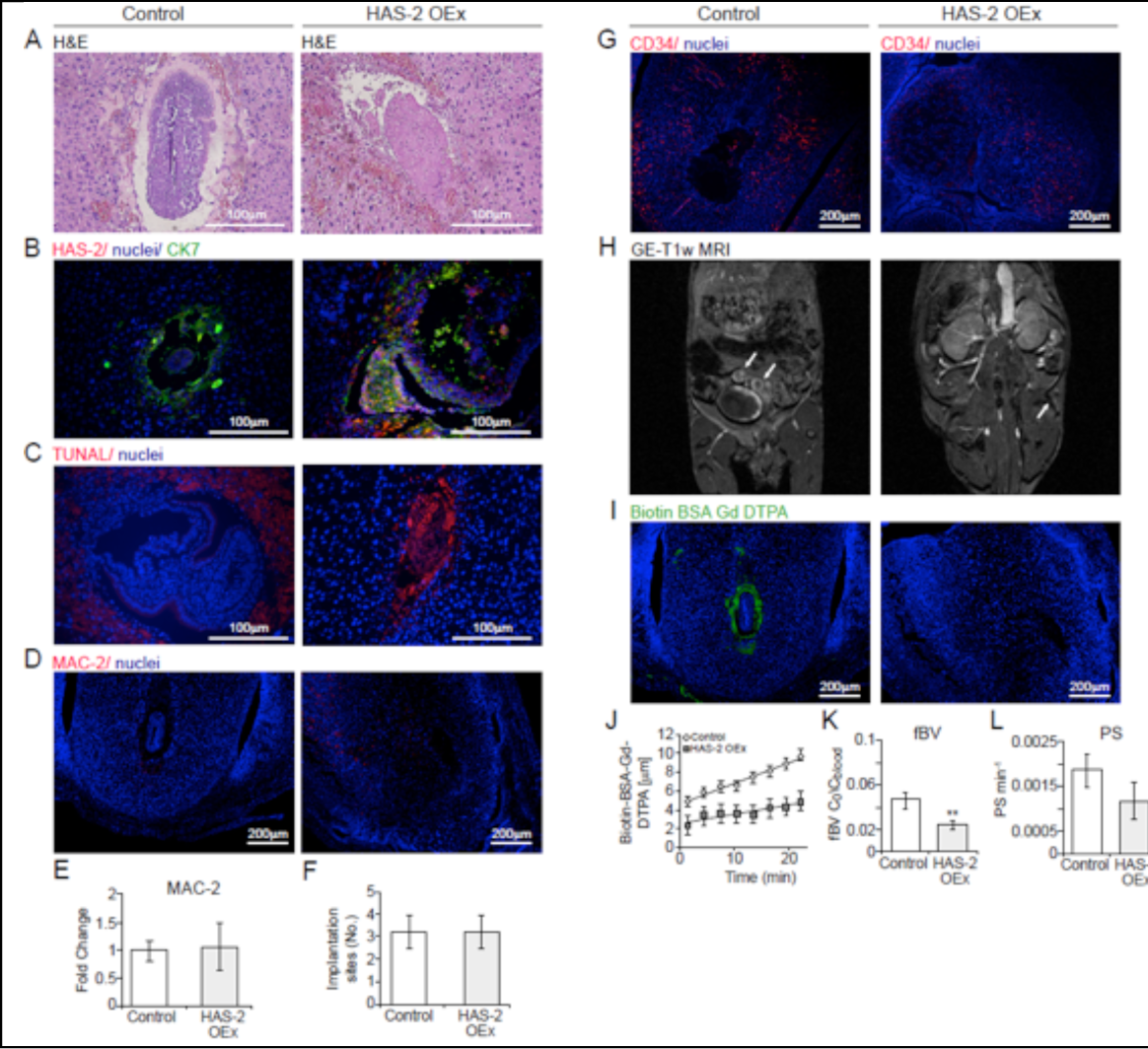
Trophectoderm over-expression of HAS-2 results in early embryonic lethality and attenuated decidual angiogenesis. Embryos over-expressing HAS-2 and eGFP or just eGFP in trophoblast cells (A-F). (**A**) H&E staining of implantation sites of embryos over-expressing HAS-2 and eGFP or eGFP alone as control, revealed embryo resorption as a result of HAS-2 OEx (n=4 dams). (**B**) Remnants of embryo over-expressing HAS-2 by trophoblast cells (CK-7) as opposed to decidual cells expressing HAS-2 in the embryonic niche in the control (white arrow, implantation site; n=3 dams). (**C**) Profound embryonic cell death was detected as a result of HAS-2 OEx, using TUNEL staining (n=3 dams). (D) Similar pattern of MAC-2^+^ macrophages observed in both groups, confined to the mesometrial pole, away from the embryonic niche (n=3 dams). (**E**) Quantification MAC-2 staining by fluorescent microscopy (1.066 fold change±0.18; 0.41; n=5 dams, 7 implantation sites, p=0.879). (**F**) Number of embryo implantation site per dam was assessed, at E6.5 by gross morphology inspection as well as by examination of histological sections (3±0.63; 2.88±0.61, p=0.88) (n=6 dams in control; 8 dams in HAS-2 OEx). (**G**) Impaired development of CD34^+^ newly formed blood vessels was observed in HAS-2 OEx pregnancies, reflected by confinement of vessels away from the embryonic niche (n=3 dams). (**H**) T1 weighted GE-MRI of embryo implantation sites, acquired from pregnant mice at E6.5 30 minutes after administration of biotin-BSA-GdDTPA. Little accumulation of biotin-BSA-GdDTPA was observed in the HAS-2 OEx group in comparison to control (white arrows, implantation sites; n= 6 dams). (**I**) Visualization of hyper-permeable blood vessels in the embryonic niche was conducted by staining of biotin-BSA-GdDTPA, 40 minutes after intravenous injection. Hyper-permeable vessels were not detected at HAS-2 OEx deciduae in contrast to those detected in the control group (n=3 dams). (**J-K**) These observations were consistent with DCE-MRI of biotin-BSA-Gd-DTPA, fBV (F; 0.041±0.005; 0.023±0.002; p= 0.004) and PS (G; 0.0017±0.0003; 0.0009±0.00031; p=0.2) (control: 5 dams 9 implantation sites; HAS-2 OEx: 6 dams 19 implantation sites).

## Discussion

During the post-implantation stage, prior to the formation of a functional hemo-chorial placenta, decidual vasculature provides the embryo with oxygen and essential nutrients by diffusion. Analyses of the embryo-maternal interface demonstrated the confinement of decidual angiogenesis to the outskirts of the implantation site in the anti-mesometrial pole, and their absence from the mesometrial pole^8^. Interestingly, this compartmentalization takes place despite the flux of paracrine, hypoxia-induced, pro-angiogenic signals secreted from uterine decidual cells adjacent to the embryo, facing both decidual poles^8^. Furthermore, the maternal primary decidual zone, at the immediate vicinity of the embryo, is characterized by impermeable vasculature. The goal of the present work was to decipher the mechanism responsible for the maintenance of a vascularpermeability barrier at the feto-maternal interface of the post-implantation embryo during the early stages of pregnancy, and further unveil its significance to pregnancy outcome.

Our experiments show that dynamic deposition and degradation of hyaluronan throughout embryo implantation, correlated, respectively, with decidual angiogenesis and vascular expansion. Lineage specific and reciprocal genetic manipulations of trophectoderm expression of key hyaluronan metabolic enzymes, provided evidence for the discrete spatio-temporal roles for hyaluronan as well as its metabolites, which are indispensable for proper development of the embryo. Specifically, our findings support the role of high molecular weight hyaluronan as a negative angiogenic morphogen during early pregnancy, further suggesting that hyaluronan breakdown products, generated upon its degradation by hyaluronidases, promote vascular permeability via the VEGF-VEGFR-2 signaling pathway, to support perfusion to the developing embryo. The temporal fine-tuning of conversion of the full-length hyaluronan into its degradation products is governed by progesterone signaling. The effect of hyaluronan in vascular remodeling and decidual homeostasis involves uterine NK cells and its contribution to trophoblast invasion is assisted by MMP-9. Schematic presentation of the inverse correlation between the spatio-temporal distribution of hyaluronan and decidual angiogenesis, resulting in vascular sub-compartmentalization, via recruitment and differentiation of uterine NK cells, is presented in **Fig 10**.

**Figure 10.**
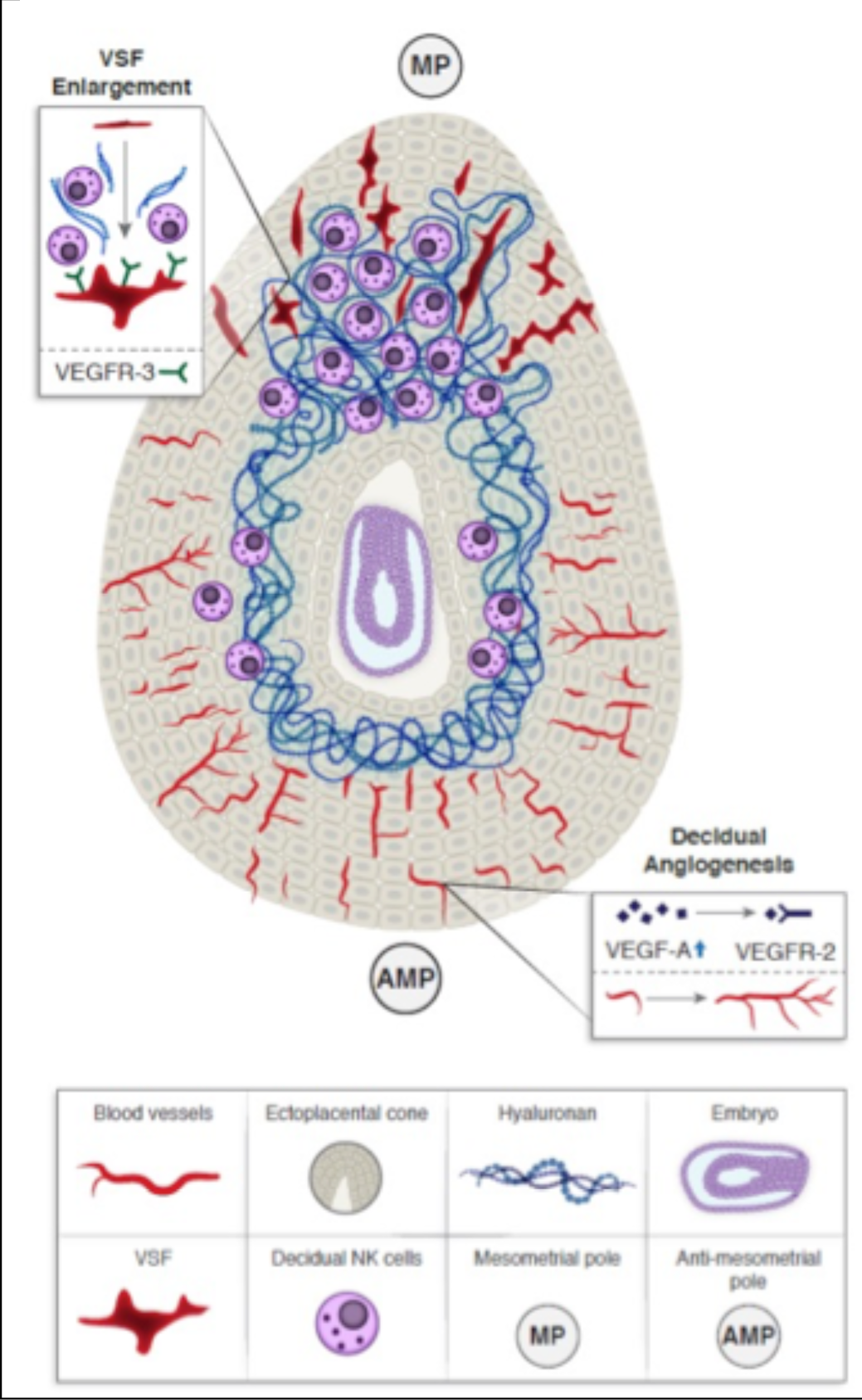
Scheme describing the regulation of hyaluronan over sub-compartmental decidual vascular remodeling via uterine NK recruitment cells during early pregnancy in mice.

To assess the involvement of progesterone in the frame of physiological decidual hyaluronan metabolism, we pharmacologically inhibited the downstream actions of its receptor, after attachment is established, in order to allow the onset of gestation. This treatment lowered the expression of Hyal-2 at E5.5 together with that of VEGF-A, previously shown as a target of decidual progesterone receptor and a regulator of the post-attachment angiogenic reaction in mice^8^. Thus, in addition to its effect on decidual angiogenesis, progesterone receptor signaling positively regulated Hyal-2 expression in primary decidual cells, enabling invasion of trophoblast cells to the embedding maternal stroma.

Expression of both hyaluronan synthesizing enzymes as well as its degrading enzymes, in the feto-maternal interface, was prominent throughout the post-implantation period. Specifically, Hyal-2 and HAS-2 was observed in giant as well as cytotrophoblast cells at E6.5.,. Subsequently to its function in hyaluronan clearance, Hyal-2, which was also observed in primary uterine decidual cells, adjacent to the embryonic niche, generates intermediate hyaluronan fragments of approximately 20 kDa, which are pro-angiogenic^27^. To study the impact of peri-embryo hyaluronan metabolism, we used lenti-viral infection generating either Hyal-2 or HAS-2 over-expressing blastocysts, targeted to their trophectoderm cells. Enhanced degradation of hyaluronan at the decidual-trophoblast interface, by Hyal-2 over-expression, resulted in smaller deciduae, cell death in the embryonic niche, as well as abnormal morphology of the embryos.

Alike connective tissue homeostasis and cancer metastasis ^33, 38^, Hyal-2 over-expression by trophoblast cells induced both, zymogen as well its mature form of MMP-9, in a similar pattern to that previously demonstrated *in vitro* ^33^. This response apparently resulted in excess of lateral and mesometrial stromal invasion at E6.5 accompanied by impaired uterine NK cells recruitment. This genetic manipulation was also accompanied by prominent ectopic angiogenesis proximal to the ecto-placental cone, manifested by local elevation in fractional blood volume and blood-vessel permeability, as well as leakage of maternal blood into the embryonic niche. The prominent increase in blood volume can be attributed to the distinctive pro-angiogenic effects of Hyal-2-mediated hyaluronan degradation products. The latter could potentially act through either the release of ECM bound VEGF by the up-regulated MMP-9 activity, as previously demonstrated in carcinogenesis ^37^, or directly via up-regulated expression of VEGF and its receptor VEGFR-2, observed herein and also demonstrated during distinct vascular remodeling events ^29, 30, 39^. Furthermore, Hyal-2 OEx resulted in compromised recruitment and function of uterine NK cells, responsible for VSF formation, via enlargement and maturation of mesometrial vasculature and prevention of vessels sprouting as previously demonstrated during post-implantation development in mice ^8, 18, 40^. Additionally, VEGFR-3 another regulator of VSF formation ^8^, has been demonstrated as an attenuator of VEGF-VEGFR-2 induced vascular permeability via inhibition of VEGFR-2 ^12^. Therefore, we cannot rule out the contribution of decreased VEGFR-3 in the mesometrial induction of VEGFR-2, resulting in vascular hyper-permeability.

Transcriptome and functional analysis of uterine NK cells, sorted from foster dams, indicated a prominent shift in uterine NK cells programming in Hyal-2 OEx surrogate mice alongside impaired recruitment and vascular remodeling attributes.

Namely, uterine NK cells displayed a phenotype similar to that of peripheral NK cells, depicted by decreased expression of vascular remodeling factors, immunomodulatory molecules and classification markers, which are usually up-regulated in uterine NK cells pre-placentation (e.g. Rgs5, Tnfrsf9, Ocel1, Gzmg, Ido3)^15, 41, 42^. Interestingly, uterine NK cells sorted from Hyal-2 OEx mice acquired an antiviral associated phenotype, exhibiting a local enhanced innate immune response. The latter might be an intermediate phenotype as a result of impaired differentiation, or an active response to excess trophoblast invasion, observed in Hyal-2 OEx deciduae.

Due to the classical role of hyaluronan as a biological glue ^22^, it has been hypothesized that it facilitates attachment of the pre-implantation embryo. This claim was supported by enhanced attachment of isolated, pre-implantation blastocysts to hyaluronan-rich matrix ^43^ and further challenged by the use of hyaluronan-enriched blastocyst transfer medium with the aim of improving blastocyst adherence ^44^. Interestingly, Hyal-2 over-expression enhanced implantation rate. This observation may reflect the potency of low-molecular-weight hyaluronan to induce cellular motility and invasion, as previously described for tumor cells, fibroblasts and endothelial cells ^34, 45–47 48^. Along this line, RHAMM, a receptor for low molecular-weight hyaluronan, which is an efficient mediator of tumor cell invasion^49^, was prominently expressed at the attachment interface. Moreover, the local induction of MMP-9 expression in the trophectoderm, as a result of Hyal-2 over-expression, may have contributed to the increased implantation rate. It has been reported that low-molecular weight hyaluronan-stimulated expression of pro-inflammatory cytokines and chemokines, triggers sterile inflammation, and activates murine pro-inflammatory M1 macrophages ^24, 50^. Indeed, Hyal-2 over-expression resulted in increased activation of macrophages in the entire decidua and, particularly, in the embryonic niche.

Augmentation of hyaluronan was induced by over-expression of HAS-2 in embryonic trophoblast cells. HAS-2 over-expression did not modify the number of embryo implantation sites, but resulted in profound embryonic cell death, at E6.5. Importantly, while over-expression of either HAS-2 or Hyal-2 brought about implantation failure, each of these treatments led to opposing vascular phenotypes. HAS-2 over-expression resulted in decreased angiogenesis, reduced blood volume fraction and attenuated permeability of maternal vessels, thus increasing the avascular niche of the implantation site and enlarging the maternal-embryo barrier. Despite the typical decidual morphology, no embryonic ’egg-cylinder’ structures were observed. These results underscore the importance of fine-tuned maternal-embryo barrier mediated by hyaluronan deposition and degradation so as to maintain embryo integrity, and assure accurate diffusion distance and spatial organization of trophoblasts, endothelial cells and immune cells.

As a whole, our study shows that hyaluronan acts as an ECM-based dynamic mediator of the primary maternal-embryo barrier for the developing embryo prior to placentation and is crucial for decidual morphogenesis. Furthermore, our findings highlight a critical role for hyaluronan as a vascular morphogen, which acts via recruitment and function of uterine NK cells, in the frame of gestational vascular adaptations. These findings implicate hyaluronan metabolism as a key regulator for the successful establishment of pregnancy.

## Methods

### Animals

C57Bl/6J female mice (6-12 week old; Envigo, Israel) were mated with Myr-Venus homozygote males ^51^. These hemizygote Myr-Venus embryos were used for histological analysis of hyaluronan metabolism. ICR males and females (8-12 week old; Envigo) were mated and assessed for the occurrence of vaginal plugs on the following day. These mice were used for gene expression analysis during early pregnancy, as well as for MRI experiments. All mice were maintained under specific pathogen–free conditions and handled under protocols approved by the Weizmann Institute Animal Care and Use Committee according to international guidelines.

*Immunohistochemistry* (see Supplementary material).

### Gene expression analysis

RNA extraction was performed according to manufacturer’s protocol (PerfectPure RNA tissue kit, 5 Prime; Gaithersburg, MD, USA) for all analyses. Then, cDNA production followed the manufacturer’s protocol with the High Capacity cDNA Reverse Transcription kit (Applied Biosystems, Foster City, CA, USA). Real-time PCR was performed to test the different HAS and Hyaluronidases isozymes and the expression of two of the hyalhedrins’ (TSG-6 and Versican) in embryo implantation sites during early pregnancy (E3.5-E6.5). Mouse B2M was used as a reference gene in all experiments. The primers used are listed in Supplementary Table 1. Real Time PCR was performed with SYBR Green (Roche, Indianapolis, IN, USA) in a Light cycler 480 machine (Roche), according to manufacturer’s protocol.

### In-vivo Dynamic Contrast–Enhanced (DCE) MRI of embryo implantation sites

MRI experiments were performed at 9.4 T on a horizontal-bore Biospec spectrometer (Bruker, Karlsruhe, Germany) using a linear coil for excitation and detection (Bruker). The animals were anesthetized with isoflurane (3% for induction, 1% to 2% for maintenance (Abbott Laboratories, Abbot Park, IL, USA) in 1 liter/min oxygen, delivered through a muzzle mask. Respiration was monitored and body temperature was maintained using a heated bed. The pregnant mice were serially scanned at E6.5 either after intra-peritoneal administration of 1µg/g BW of hyaluronan synthesis inhibitor 6-diazo-5-oxo-1-norleucine (DON, Sigma-Aldrich) at E3.5-E5.5, or after transgenic embryo transplantation. Three-dimensional gradient echo (3D-GE) images of the implantation sites were acquired before, and sequentially, for 40 minutes after intravenous administration of contrast agent. A series of variable flip angle, pre-contrast T_1_-weighted 3D-GE images were acquired to determine the pre-contrast R_1_ (repetition time (TR): 10 msec; echo time (TE): 2.8 msec; flip angles: 5°, 15°, 30°, 50°, 70°; two averages; matrix: 256 × 256 × 64; field of view (FOV): 35 × 35 × 35 mm^3^). Post-contrast images were obtained with a single flip angle (15°). During MRI experiments, the macromolecular contrast agent biotin-BSA-GdDTPA (80 kDa; Symo-Chem, Eindhoven, The Netherlands), 10 mg/mouse in 0.2 mL of PBS, was injected intravenously through a pre-placed silicone catheter inserted into the tail vein.

The MRI scans allowed quantification of the fBV and the PS of embryo implantation sites, as previously reported ^9^. In brief, the change in the concentration of the administered biotin-BSA-GdDTPA over time (*Ct*), in the region of interest, was divided by its concentration in blood (*Cblood*; calculated in the region of interest depicting the vena cava, also acquired during MRI, and extrapolated to time 0). Linear regression of these temporal changes in *Ct*/*Cblood* yielded two parameters that characterize vascular development and function: (i) fractional Blood Volume (fBV = *C0*/*Cblood*), which describes blood-vessel density and is derived from the extrapolated concentration of the contrast agent implantation sites, at time zero, divided by the measured concentration in the vena cava, approximately 5 minutes after i.v administration, and (ii) Permeability Surface area product, *PS* [= (*Ct – C0*)/(*Cblood* × *t*)], which represents the rate of contrast agent extravasation from blood vessels and its accumulation in the interstitial space, and is derived from the slope of the linear regression of the first 15 minutes after contrast agent administration (*t* = 15). Mean fBV and PS were calculated separately for single implantation sites considering homogeneity of variances between mice.

At the end of the MRI session, embryo implantation sites were harvested and immediately placed in 4% PFA after sacrificing the pregnant mice by cervical dislocation.

### Scanning electron microscopy

Pregnant mice were sacrificed at E6.5. Their uterine horns were harvested and fixated overnight with Kranovsky fixative -4% PFA, 2% glutaraldehyde (GA) in 0.1 M Cacodilate (Caco) buffer. After fixation, tissues were dissected and embedded into 9% Agarose gel in needed orientation, then cut on vibrotome (EMS) into 300 µm thickness slices. The slices were then stained with 1% OsO_4_ in Caco 0.1 M buffer, dehydrated in ethanol series, (50%, 70%, 96%, and 100%), and then dried using critical-point dryer BAL-TEC CPD 030. The dry samples were placed on aluminum stabs covered with carbon tape and then coated with thin layer of gold/palladium alloy in EDWARDS sputter coater. The samples were visualized using secondary electron (SE) detector in a high-resolution Ultra 55 SEM (Zeiss).

### Progesterone receptor pharmacological blockade

Pregnant mice (late E4.5) were administered intraperitoneally with RU486 (8mg/kg, Sigma-Aldrich), sacrificed at E5.5 and their uteri examined via histological analyses.

### Generation of transgenic trophoblasts in murine blastocysts. Lentiviral vectors design and production

Lentiviral vectors were constructed to produce lentiviruses expressing mouse *Hyaluronidase-2* (Hyal-2), and mouse *Hyaluronan synthase-2* (HAS-2). Mouse *Hyal-2* (GeneBank accession No. <u>NM_010489.2</u>) was isolated from uterine cDNA by PCR using a sense primer carrying a Human influenza hemagglutinin (HA) tag and restriction sites for AgeI and the anti-sense primer for SfiI (Supplementary Table 2).

Mouse *HAS-2* (GeneBank accession No. <u>NM_008216.3</u>) was isolated from uterine cDNA by restriction-free cloning, using primers containing complementary overhangs to the designated target vector (Supplementary Table 2). The purified PCR products were cloned into the lentiviral expression vector, pCSC-SP-PW-IRES/GFP (kindly provided by Professor Alon Chen, Department of Neurobiology, The Weizmann Institute of Science, Rehovot, Israel). Recombinant lentiviruses were produced by transient transfection in HEK293FT cells (Invitrogen, CA, USA), as described earlier ^52^, using 3 envelope and packaging plasmids and one of three viral constructs: (i) pCSC-SP-PW-Hyal-2-IRES/eGFP (Hyal-2 OEx), (ii) pCSC-SP-PW-HAS-2-IRES/eGFP (HAS-2 OEx), or (iii) pCSC-SP-PW-IRES/eGFP (control). Briefly, infectious lentiviruses were harvested at 48 and 72 hours post-transfection, filtered through 0.45-mm-pore cellulose acetate filters and concentrated by ultracentrifugation at 19400rpm, 4^0^C for 2.5 hours.. Lentiviral supernatant titers were determined by Lenti-X p24 Rapid Titer Kit (supplemental Table 3) according to manufacturer’s protocol (Takara Bio USA, Inc. California, U.S.A).

### Validation of gene over-expression

ES-2 cells, ovarian clear cell carcinoma cell line, for generation of lentiviruses, were used for validation of both Hyal-2 and HAS-2 over-expression after infection with lentiviral vectors. Validation was conducted using Western blot analysis (with the antibodies used for immunohistochemistry). Blastocysts from all three groups were stained by whole-mount incubation with antibodies against Hyal-2 and HAS-2, stained with species specific secondary antibodies, counterstained with Hoechst (Invitrogen), subsequently mounted in mineral oil, imaged using spinning disk 386 confocal microscope as well as 710 confocal microscope (Zeiss, Cell observer SD.) and quantified by ImageJ software (https://imagej.nih.gov/ij/).

### Mice and lentiviral transduction

Wild-type ICR females were super-ovulated by sub-cutaneous injection of pregnant mare’s serum gonadotropin (PMSG) (Sigma-Aldrich) (5 units) followed 48 hours later by intra-peritoneal injection of human chorionic gonadotropin (hCG) (Sigma-Aldrich) (5 units) and then mated with wild-type ICR males. Morulae stage embryos were collected from the females at E2.5 and then incubated in KSOM medium to obtain expanded blastocysts. Zona pellucida was removed in acidic Tyrode’s solution ^53^. Next, 15-30 embryos were incubated with lentiviruses, described above, in KSOM for 5 hours. The transduced blastocysts were washed four times, and then implanted into pseudopregnant ICR females generated after mating with vasectomized ICR males. We transplanted ∼ 10 blastocysts into each mouse using Non-Surgical Embryo Transfer kit [NSET] (ParaTechs, Lexington, KY, USA). Imaging of infected placenta was performed using a fluorescent Olympus SZX-RFL2 zoom stereo microscope.

### Flow cytometry analysis

Cells were isolated from deciduae harvested from foster dams at E6.5, subjected to mechanical fragmentation and followed by enzymatic digestion with 1 mg/ml collagenase type IV (Sigma-Aldrich), 0.2 mg/ml DNase (Roche) in PBS with MgCl2 and CaCl2 (Sigma-Aldrich) in 37°C for 30 minutes. Then cells were stained with PE coupled antibody to mouse NCR1 (29A1.4, BioLegend, CA, USA) and APC coupled antibody to mouse CD45 (30-F11, BioLegend). Cells were then passed through a 70µm mesh and washed in FACS buffer (2Mm EDTA, 0.5% BSA in PBS). Cells were then subjected to flow cytometry by a SORP-FACSAriaII machine using a 70 µm nozzle, and analyzed by FACSDiva 8.0.1 software (BD).

### Transcriptomics sequencing

Massively Parallel Single-Cell RNA-seq library preparation (MARS-seq) NK cell Libraries were prepared as previously described^54^. In brief, 20000 NCR1+ cells were sorted into 40µl of lysis/binding buffer, from which, mRNA was captured with 12 ml of Dynabeads oligo(dT) (Dynabeads™ mRNA DIRECT™ Purification Kit, NY, USA), washed, and eluted at 85C with 10 ml of 10 mM Tris-Cl (pH 7.5) and processed according to protocol previously developed for single-cell RNA-seq^54^. In brief, the samples were barcoded, converted to cDNA, pooled and linearly amplified by T7 *in vitro* transcription. mRNA was then fragmented and converted into a sequencing-ready library, which is then tested for quality and concentration^54^. MARS-seq libraries were sequenced using an Illumina NextSeq 500 sequencer, at a sequencing depth of ∼ 5 million reads per sample.

### Statistics

All statistical analyses used in this study were two-tailed with a similar level of significance (p=0.05), and demonstrated normal values distribution. Beside the on-way ANOVA analysis conducted for gene expression analysis of hyaluronan metabolism during implantation (**Figure S1**), all statistical analysis were examined by *t* tests, while assuming homogenous distribution of variances. All statistical analyses were conducted using GraphPad Prism 7(CA, USA).

### Bioinformatic analysis

MARS-seq analysis was done using the UTAP transcriptome analysis pipeline^55^ at the bioinformatics unit. Reads were trimmed using cutadapt (DOI: 10.14806/ej.17.1.200) and mapped to the Mus_musculus genome (UCSC mm10) using STAR (DOI: 10.1093/bioinformatics/bts635) v2.4.2a with default parameters. The pipeline quantifies the genes annotated in RefSeq (that have expanded with 1000 bases toward 5’ edge and 100 bases toward 3’ bases). Counting was done using htseq-count (DOI: 10.1093/bioinformatics/btu638) (union mode). Genes having minimum 5 UMI-corrected reads in at least one sample, were considered. Normalization of the counts and differential expression analysis was performed using DESeq2 (DOI: 10.1186/s13059-014-0550-8) with the parameters: betaPrior=True, cooksCutoff=FALSE, independentFiltering=FALSE. Raw P-values were adjusted for multiple testing using the procedure of Benjamini and Hochberg.

Functional analysis of differentially expressed genes (FDR < 0.05) was conducted using GeneAnalytics (https://ga.genecards.org) with default settings. The Holm-Bonferroni test correction was implemented to detect pathway enrichment and GO: Molecular Functions analyses. Gene Set Enrichment Analysis (GSEA) was performed using GSEA 3.0 with the GSEAPreranked tool (reference: Subramanian, Tamayo, et al. (2005, PNAS 102, 15545-15550) and Mootha, Lindgren, et al. (2003, Nat Genet 34, 267-273). Gene names were converted to human gene symbols, and analyzed with default parameters. The Molecular Signature Database, with Biological Processes and Hallmark gene sets, were used to perform pathway enrichment analysis. Gene expression levels (rld) of specific leading edge subsets were visualized as heatmaps using the Partek software.

*ELISA.* Embryo implantation sites were harvested from foster dams at E6.5. ELISA was performed on total extracted protein according to manufacturer instructions for Mouse IL-15/IL-15R Complex (NBP1-92667, Novus Biologicals, CO, USA). Results represent five pools of three tissues harvested from each foster dam.

## Author contributions

R.H designed research studies, conducted experiments, acquired data, analyzed data, and wrote the manuscript. E.G designed research studies and conducted experiments. A.C designed research studies, conducted experiments, acquired data and analyzed data. S.S conducted experiments, acquired data and analyzed data. O.A conducted experiments and acquired data. S.L designed research studies, conducted experiments and acquired data. O.G analyzed data. M.E conducted experiments. G.C conducted experiments. E.K conducted experiments and acquired data. R.E conducted experiments, acquired data and analyzed data. N.D designed research studies and wrote the manuscript. M.N designed research studies and wrote the manuscript.

## Acknowledgement

We thank Prof. Karina Yaniv, Prof. Joel Garbow and Dr. Tal Raz and Prof. Alon Chen for useful discussions. RNAseq work was done with critical advice from Dr. Hadas Keren-Shaul from the Sandbox unit in ‘Life Science Core Facility of Weizmann Institute of Science.’

This work was supported by the Seventh Framework European Research Council Advanced Grant 232640-IMAGO and by National Institutes of Health (grant 1R01HD086323-01). Michal Neeman is incumbent of the Helen and Morris Mauerberger Chair in Biological Sciences.

## Supplemental Inventory

Supplementary materials and methods

Table S1-S3

**Figure S1. Hyaluronan metabolism following implantation.** Related to figure 2.

**Figure S2. Hyal-2 over-expression modified uterine vascular remodeling and enhanced trophoblast decidual invasion.** Related to figures 5-7.

