## supplementary figures for "Hyaluronan-NK cell Interaction Controls the Primary Vascular Barrier during Early Pregnancy"

### Supplementary Methods

*Immunohistochemistry*. Uterine sections containing embryo implantation sites were fixed in 4% paraformaldehyde (PFA) and embedded in paraffin. For morphological analysis, tissues were stained with hematoxylin and eosin, whereas consecutive sections underwent immunohistochemical staining. The latter included antigen retrieval in citrate (pH=6.0) or EDTA (pH=8.0) buffers in a pressure cooker at 125ºC for 3 minutes. Antigens were washed, blocked with 20% normal horse serum, permeabilized with 0.2% Triton X-100 in PBS, for 1.5 hours at RT, then incubated overnight with the following primary antibodies: goat anti HAS-1 (sc-23145; Santa Cruz Biotechnology, Dallas, TX, USA) goat anti HAS-2 (sc-34067; Santa Cruz Biotechnology), rabbit anti Hyal-2 (ab68608; Abcam, Cambridge, UK), rabbit anti Hyal-1 (ab203293; Abcam) goat anti GFP (ab6673; Abcam), rabbit anti CD44 (ab41478; Abcam), rabbit anti LYVE-1 (70R-LR003; Fitzgerald Industries International, Acton, MA, USA), rat anti CD34 (CL8927PE; Cedarlane, Ontario, Canada), Rabbit anti MMP-9 (ab38898; Abcam), Rabbit anti Cytokeratin-8 (NB110-56919; Novus biologicals, CO, USA), Mouse anti Cytokeratin-7 (ab9021;Abcam) and rat anti-mac-2 (CL8942AP; Cedarlane), rat anti mouse VEGFR-3 (14-5988-81; eBioscience, MA, USA),

Rabbit anti VEGFR-2 (CST-9698S, Cell Signaling, MA, USA) and biotin conjugated DBA (L2785; Sigma-Aldrich, Rehovot, Israel) (ab199128; Abcam, Cambridge, UK). Next, slides were washed and incubated with secondary antibodies, conjugated to biotin against the appropriate species (except for goat anti GFP in the case of double staining, diluted 1:100 in 2% normal horse serum in PBS, for 1.5 hours at RT. Slides were then washed in PBS and incubated in Cy3 or Cy2-conjugated streptAvidin (Jackson Immunoresearch Laboratories, PA, USA), diluted 1:150 in PBS for 45 minutes, at RT. Cells undergoing apoptosis were detected by TUNEL staining (ApopTag; Merck Millipore, MA USA). Slides were counterstained with Hoechst (Invitrogen, Carlsbad, CA, USA) and subsequently mounted. For staining of hyaluronan, sFlt-1 or VEGF-A, sections were bleached by blocking endogenous peroxidase activity, 1% $H_2O_2$ in methanol, for 30 minutes. Then, antigen retrieval in citrate (pH=6.0) in a pressure cooker at 125ºC for 3 minutes, washed, blocked with 20% normal horse serum, permeabilized with 0.05% Tween-20 in PBS, for 1.5 hours at RT, followed by incubation with sheep anti hyaluronan antibody, Rabbit anti sFlt-1 or Rabbit anti VEGF-A (ab53842, Abcam; 36-1100, Invitrogen; sc-152-G, Santa Cruz respectively) for 2 days in 4°C in a wet chamber. Next, slides were washed and incubated with the appropriate secondary antibodies, diluted 1:100 in 2% normal horse serum in PBS, for 1.5 hours at RT. Then, slides were washed in PBS and incubated in avidin/biotin complex (ABC, Vectastatin, ABC, Vector Labs, CA, USA) in PBS, for 1.5 hours at RT, followed by exposure to DAB (Sigma-Aldrich, Rehovot, Israel). Slides were counterstained with hematoxylin and subsequently mounted. All slides were imaged using a fluorescent Olympus SZX-RFL2 zoom stereo microscope.

*Ex-vivo separation of glycosaminoglycans and characterization of HA fragments by gel electrophoresis*. Uterine glycoseaminoglycans separation was carried out as follows: A pool of 3 implantation sites per time-point, was lyophilized overnight. Wet/dry weight was determined by weighing the samples before and after lyophilization. Samples were digested in 950µl 0.0005% Phenol Red, 100 mM ammonium acetate, ph 7.0 containing 125 mg proteinase K (Roche) for two hours at 60 °C Another 125 mg of proteinase K was added to each tube and incubated for another two hours. The samples were boiled to inactivate the proteinase K and pelleted by centrifugation to remove any undigested material. Samples were subjected to DNA (20U,DNAse1, Roche) and RNA degradation (192U,RNAse A, Roche), followed by enzyme heat inactivation. Glycoseaminoglycans were precipitated in Ethanol overnight at -20°C, pelleted by centrifugation, re-suspended in 20 µl (Tris/Borate/EDTA [TBE]1x with loading buffer (TBE, 0.02% bromophenol blue and 2M sucrose) to a total of 25 µl. Select-HA LoLadder (Ambion, UK) was used as molecular weight gel standard. Hyaluronan was separated on Tris-glycine Native gels (Any-kD polyacrylamid gel, $8.6^x 6.7$ cm ($W^x L$) (BIO-RAD, Israel)).

Outer and inner buffer chambers were filled with pre-cooled TBE. Gels were run at 300 V for 1 hour, rinsed in water, then transferred to staining solution (0.005% Stains-All (Sigma), 50% ethanol, 50% water) in the dark overnight. Gels were de-stained in water and imaged on a flatbed scanner.

*Western blot analysis.* Pools of two deciduae per mouse were harvested from Control and Hyal-2 OEx foster Dams. Cells were lysed in RIPA buffer and equal amounts of protein were separated by either 12 % or 8% SDS-PAGE and transferred to nitrocellulose membranes. For detection, we used Rabbit anti MMP-9 (ab38898; Abcam), Rat anti VEGFR-1 (MAB471; R&D, MN, USA), Rabbit anti VEGF (ab52917; Abcam), Rabbit anti VEGFR-2 (55B11; Cell signaling, MA, USA), Rat anti mouse VEGFR-3 (14-5988-81; eBioscience, MA, USA), Goat anti NCR-1 (AF2225; R&D, MN, USA) and Rabbit anti GAPDH (C-2118S; Cell signaling) antibodies.

*Quantification of trophoblast invasion.* Post-implantation invading trophoblasts were detected by immunolabeling of cytokeratin-8 (CK-8) on paraffin embedded histological section retrieved from Control and Hyal-2 OEx foster Dams. Invading cells were quantified within several rings around the core of the egg-cylinder, from the CK-8 positive visceral endoderm cells, outward, to the decidual stroma. Analysis and final quantification were performed using ImageJ software (https://imagej.nih.gov/ij/), excluding auto-fluorescent erythrocytes detected by DNA Hoechst labeling.

*Clearing and immunolabeling.* Surrogate mothers carrying both Hyal-2 OEx and control embryos, were i.v administered with 10 mg/kg ROX (Molecular Probes) conjugated Lycopersicon Esculentum lectin (Vector Laboratories) for detection of functional blood vessels. Ten minutes after administration, mice were euthanized with pentobarbital (CTS Chemical Industries) and were immediately perfused with 1% heparin (heparin sodium Fresenius, Bodene (PTY) limited) in PBS followed by 4% PFA. Embryo implantation sites were harvested, fixed with 4% PFA for 3 days at 4°C, washed three times with PBS, permeabilized with [0.2% Triton X-100(TX-100] for 4h, and placed overnight in blocking solution [10% normal horse serum in PBS containing 0.05% TX-100], incubated in 1:400 dilution of primary rat anti mouse MAC-2 antibody, and 1:200 dilution of rabbit anti GFP antibody (ab6556, Abcam) in antibody cocktail (50% blocking solution/0.05% TX-100/PBSX1) for 1 week at 4°C, washed for 24h with washing buffer (1% blocking solution/0.05% TX-100/PBS, and incubated with a mixture of secondary Cy5 (1:250 dilution, Jackson Immunoresearch Laboratories) in antibody cocktail for another week at 4°C. Following the 24h wash, samples were subjected to a clearing procedure. For tissue clearing, we used a modified protocol based on Chung et al., 2013 and Hama et al., 2011. Briefly, following staining, tissues were removed to hydrogel solution (4% Acrylamide, 0.025% Bisacrylamide, 0.25% Va-

044, 4% PFA in PBSX1) for 2 weeks followed by passive clearing process (200mM Boric acid, 4% SDS) for 2 weeks at 37°C. In the next step, samples were placed in ScaleA2 (4M Urea, 10% Glycerol, 0.1% TX-100) for 36h. 3D images and movies were acquired using a Zeiss LSM710 confocal microscope (Zeiss, Oberkochen, Germany).

### Supplemental figures

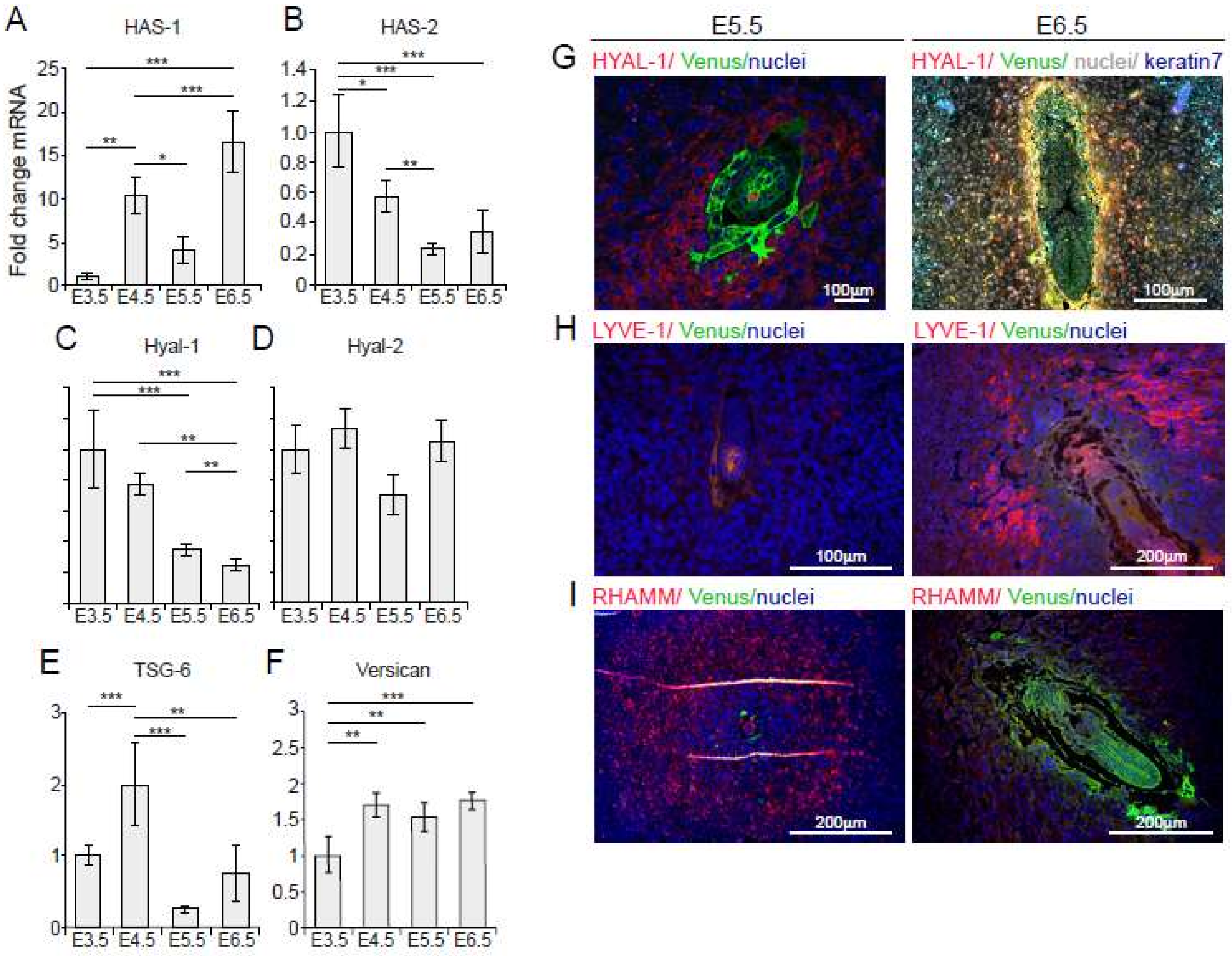


Figure S1. **Hyaluronan metabolism following implantation.** (**A-F**) Equal number of embryo implantation sites were harvested from each pregnant mouse. A total of 4 mice was examined at different time points. RNA was extracted and subjected to real-time PCR analysis. Histological analysis of post-implantation deciduae (E5.5-E6.5) (n=4). (**G**) Distribution of Hyal-1, hyaluronan degrading enzyme, during implantation. (**H**) Representative image of hyaluronan receptor, LYVE-1 decidual distribution, during implantation (n=3 dams). (**I**) Representative image of hyaluronan receptor, RHAMM distribution during implantation in (n=3 dams).

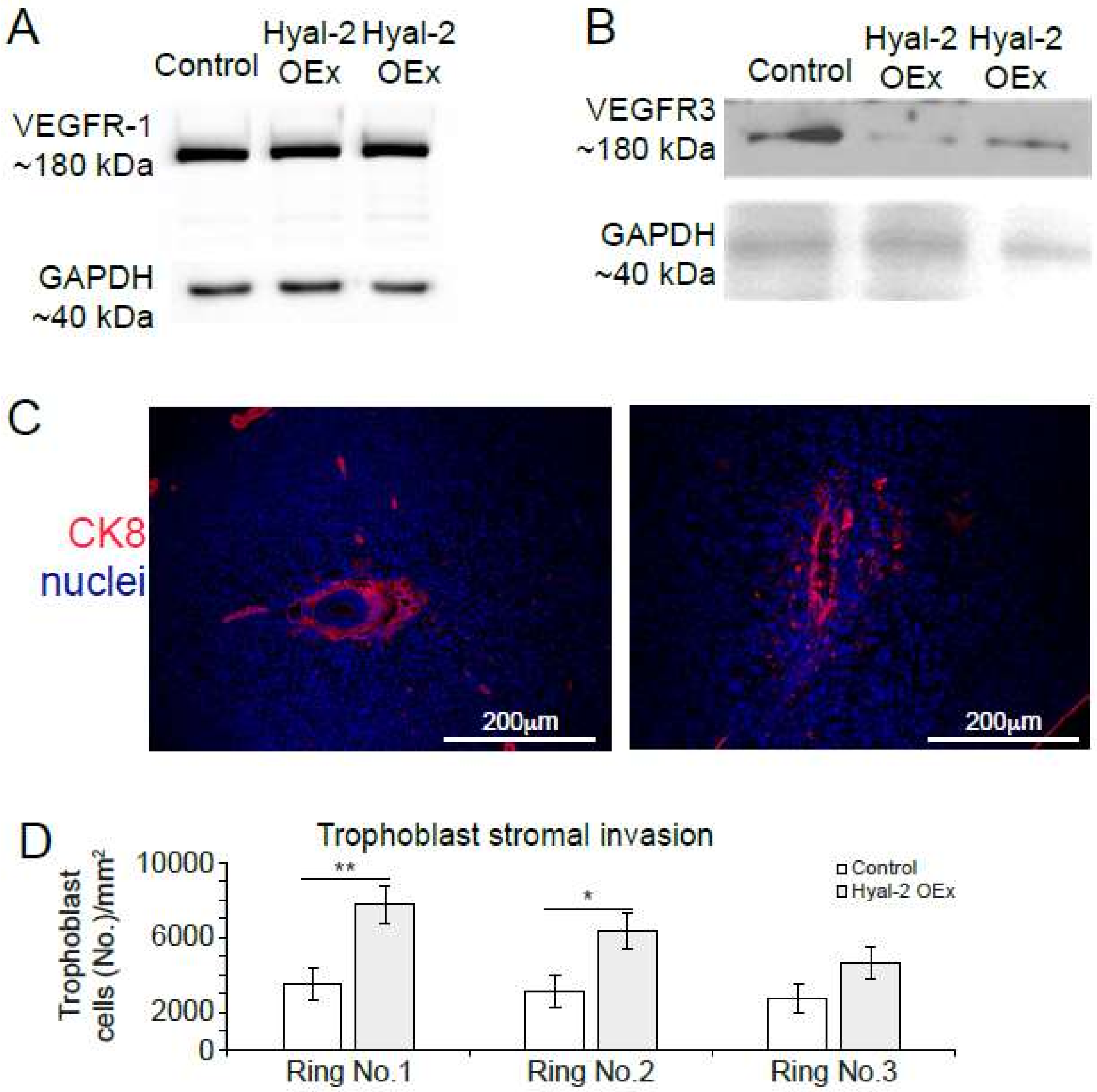


Figure S2. **Hyal-2 over-expression results increased MMP-9 levels and enhanced trophoblast decidual invasion.** (**A**) Similar levels of VEGFR-1 in deciduae harvested at E6.5 (n=3 dams 2 implantation sites from each group) (**B**) Elevated expression of VEGFR-3 in deciduae harvested at E6.5 (n=3 dams 2 implantation sites from each group). (**C-D**) Excessive invasion of trophoblast cells detected in Hyal-2 over-expressing trophoblast cells (CK-8) in decidual stroma surrounding the embryo.

**Table S1.** Real Time PCR primers

| **Gene** | **Forward primer** | **Reverse primer** |
|---|---|---|
| Mouse *HAS*-1 | CTACGTGCAGGTCTGTGACTC | GCTCGTTCCACATTGAAGGC |
| Mouse *HAS-2* | GGCCGGTCGTCTCAAATTCA | ACAATGCATCTTGTTCAGCTCC |
| Mouse *Hyal*-1 | TTCAGTCCTGAGGTTTCCCCA | GGTTGGATACCACGGAACCT |
| Mouse *Hyal*-2 | CTCAGCTGGCGTCATCTTCT | GCCCAGGACACGTTGACTAT |
| Mouse TSG-6 (*Tnfaip6*) | AGTGAGCGATGGGATGCCTAT | TCGCTTCGGATCTGTGAAGA |
| Mouse *Versican* (V-1) | CGTTTTGAGAACCAGACATGC TT | TTGGATGACCACTTACAATCA TATCA |
| Mouse *B2M* | CCCGCCTCACATTGAAATCC | GCGTATGTATCAGTCTCAGTG G |

**Table S2.** Primers used for cloning

| Vector | Insert | Forward primer | Reverse primer |
|---|---|---|---|
| pCSC-SP-PW-IRES/eGFP | HA-Hyal-2 | ATATACCGGTGCCATGGCTTACCATACGATGTTCCAGATTACATGCGGGCAGGACTAGGTCC | ATATGGCCGCCTGGGCCTCATAAGGTCCAGGTGAGAG |
| | HAS-2 | AGAAGACACCGACTCTAGAGGATCCATGCATTGTGAGAGGTTTCTATGT | GGGGGGGGCGGAATTCTGCAGTCATACATCAAGCACCATGTCA |

**Table S3.** Viral titers measured for lentiviral vectors

| Insert | Vector | Viral titer (IFU/ml) |
|---|---|---|
| Control | pCSC_SPPWCMV_IRES_GFP | ≥ 1 x $10^7$ |
| Hyal-2 CDS | pCSC_SPPW_CMV_Hyal-2_IRES_GFP | ≥1 x $10^7$ |
| HAS-2 CDS | pCSC_SPPW_CMV_HAS-2_IRES_GFP | ≥1 x $10^7$ |
